# Dermal appendage-dependent patterning of zebrafish *atoh1a+* Merkel cells

**DOI:** 10.1101/2022.08.11.503570

**Authors:** Tanya L. Brown, Emma C. Horton, Evan W. Craig, Madeleine N. Hewitt, Nathaniel G. Yee, Camille E.A. Goo, Everett T. Fan, Erik C. Black, David W. Raible, Jeffrey P. Rasmussen

## Abstract

Touch system function requires precise interactions between specialized skin cells and somatosensory axons, as exemplified by the vertebrate mechanosensory Merkel cell-neurite complex. Development and patterning of Merkel cells and associated neurites during skin organogenesis remains poorly understood, partly due to the *in utero* development of mammalian embryos. Here, we discover Merkel cells in the zebrafish epidermis and identify Atonal homolog 1a (Atoh1a) as a marker of zebrafish Merkel cells. We show that zebrafish Merkel cells derive from basal keratinocytes, express neurosecretory and mechanosensory machinery, extend actin-rich microvilli, and complex with somatosensory axons, all hallmarks of mammalian Merkel cells. Merkel cells populate all major adult skin compartments, with region-specific densities and distribution patterns. *In vivo* photoconversion reveals that Merkel cells undergo steady loss and replenishment during skin homeostasis. Merkel cells develop concomitant with dermal appendages along the trunk, and preventing dermal appendage formation reduces Merkel cell density by affecting both cell differentiation and maintenance. By contrast, altering dermal appendage morphology changes the distribution, but not density, of Merkel cells. Overall, our studies provide insights into touch system maturation during skin organogenesis and establish zebrafish as an experimentally accessible *in vivo* model for the study of Merkel cell biology.

## INTRODUCTION

Skin functions as our primary interface with the physical environment and can distinguish a range of tactile inputs with exquisite acuity. As the skin undergoes organogenesis, the epidermis transforms from a simple, uniform epithelium into a complex, diverse tissue. During these dramatic changes, the skin develops regionally specialized sensory structures and becomes innervated by specific types of somatosensory neurites (reviewed by Jenkins and Lumpkin, 2017). Interactions between somatosensory neurites and cutaneous cell types regulate diverse tactile responses (reviewed by Handler and Ginty, 2021). Altered tactile sensitivity during early mammalian development has been associated with neurodevelopmental disorders (reviewed by Orefice, 2020), underscoring the importance of understanding the cellular and molecular basis of touch system development and function.

Merkel cells (MCs), a specialized mechanosensory cell type found in the vertebrate epidermis (reviewed by Hartschuh et al., 1986), densely populate many highly sensitive regions of skin (Lacour et al., 1991). MCs have several defining cellular characteristics that distinguish them from other epidermal cell types: they are relatively small, extend actin-rich microvilli, contain cytoplasmic granules reminiscent of synaptic vesicles, and form contacts with somatosensory axons (Hartschuh and Weihe, 1980; Mihara et al., 1979; Smith, Jr, 1977; Toyoshima et al., 1998). In mammals, a subset of cutaneous somatosensory axons known as Aß slowly adapting type I low-threshold mechanoreceptors (SAI-LTMRs) innervate MCs, forming the MC-neurite complex. MCs detect mechanical inputs via the cation channel Piezo2 (Ikeda et al., 2014; Maksimovic et al., 2014; Woo et al., 2014) and play an active role in touch sensation by releasing neurotransmitters to activate neighboring neurites (Chang et al., 2016; Chang and Gu, 2020; Hoffman et al., 2018). Genetic ablation of rodent MCs indicates they are required for specific aspects of touch system function, including promoting the static phase of the slowly adapting response of Aß SAI-LTMRs and sensory tasks such as texture discrimination (Maricich et al., 2012, 2009).

Molecular control of MC development has primarily been studied in rodent hairy skin (reviewed by Oss-Ronen and Cohen, 2021). While this system has been useful for understanding many aspects of MC development and function, it also has several significant limitations. First, vertebrates have diverse types of skin and MCs are found in both hairy and glabrous (non-hairy) skin, as well as mucocutaneous regions such as the gingiva and palate (Lacour et al., 1991; Moayedi et al., 2021). Importantly, MC populations within different skin compartments share similar transcriptional profiles (Nguyen et al., 2019). Thus, the establishment of complementary genetic systems in different types of skin could help reveal both shared and divergent principles of MC development. Second, because *in utero* development of mammalian skin limits access to the developing touch system—combined with technical limitations of imaging intact mammalian skin—the dynamics of MC development and innervation remain essentially unknown. Third, unbiased screens for regulators of MC development would be difficult or impractical in rodents due to the prohibitive cost of animal housing and difficulty of visualizing MCs *in situ*.

Anamniote model systems, such as the genetically tractable zebrafish, provide the potential to overcome these limitations. Interestingly, despite the different tactile environments encountered by terrestrial and aquatic vertebrates, MCs have been described by transmission electron microscopy (TEM) in a wide variety of anamniotes, including teleost (ray-finned) fish, lungfish, and lamprey (Fox et al., 1980; Lane and Whitear, 1977; Whitear and Lane, 1981). Here, we identify and characterize a population of zebrafish epidermal cells that we propose are bona fide MCs. Our studies establish the zebrafish as a promising new model to investigate the developmental and cell biology of MCs.

## RESULTS

### Ultrastructural identification of presumptive MCs in the adult epidermis

Given the presence of cells with the ultrastructural characteristics of MCs in several teleosts (Lane and Whitear, 1977), we reasoned that the zebrafish epidermis may contain similar cells. Whitear (1989) defined five ultrastructural criteria for the identification of vertebrate MCs: 1) a relatively small volume of cytoplasm; 2) an association with a nerve fiber; 3) the presence of cytoplasmic granules; 4) desmosomal attachments to neighboring cells; and 5) peripheral microvilli.

We previously demonstrated that somatosensory axons densely innervate the epidermis above scales (Rasmussen et al., 2018), dermal appendages that cover the adult zebrafish trunk (Figure 1A). By TEM, we found that many of the axon endings in the scale epidermis arborize between keratinocyte membranes (Rasmussen et al., 2018). Interestingly, however, we identified additional axon-associated epidermal cells that were distinct from the large, cuboidal keratinocytes that comprise most of the epidermis based on several characteristics. The cells appeared relatively small and spherical with a low cytoplasmic-to-nuclear ratio (Figure 1B,C), contained cytoplasmic vesicles that in some instances localized adjacent to axon contacts (Figure 1B’) and formed desmosomal-like attachments with neighboring keratinocytes (Figure 1B’’,B’’’). Furthermore, the cells extended spike-like microvillar processes that contacted adjacent cells (Figure 1C,C’). Thus, based on established TEM criteria, we identified presumptive MCs in the adult scale epidermis.

**Figure 1.**
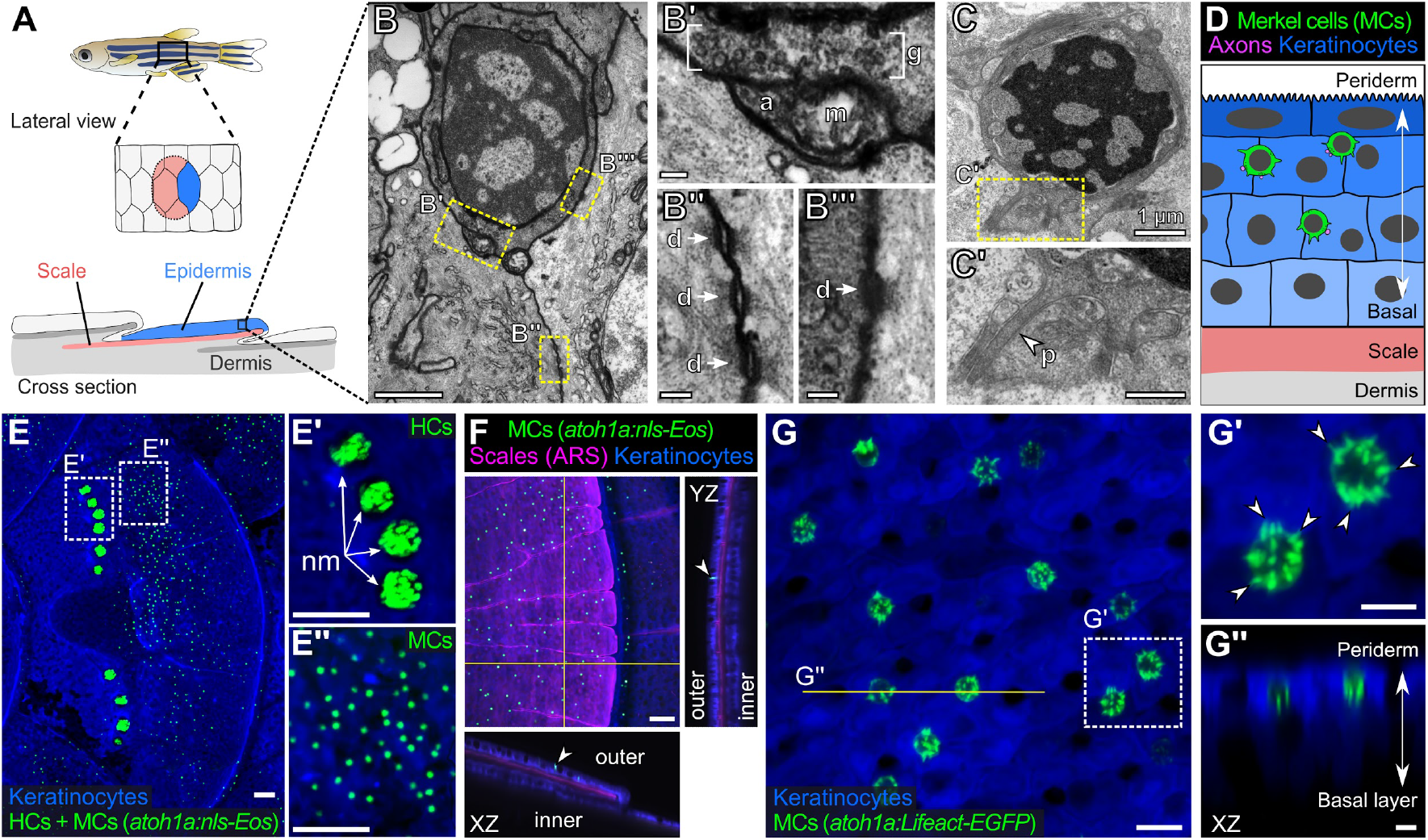
The adult scale epidermis contains *atoh1a*+ Merkel cells. Illustration of the adult zebrafish trunk anatomy showing the organization of epidermis, scales, and dermis. Scales are flat bony discs arranged in an overlapping, imbricated pattern and coated on their external surface by epidermis. **(B)** TEM of a presumptive MC from the scale epidermis. Dotted boxes indicate regions of magnification in B’-B’’’. **(B’)** Magnification of B showing cytoplasmic granules (g, brackets) juxtaposed to a putative axon (a) contact containing a mitochondrion (m). **(B’’, B’’’)** Magnifications of B showing desmosomal-like (d, arrows) attachments between keratinocytes (B’’) and between a presumptive MC and keratinocyte (B’’’). **(C, C’)** TEM of a presumptive MC from the scale epidermis showing a microvillar process (p, arrowhead). **(D)** Illustration of a cross section of the scale epidermis based on TEM observations. Superficial periderm cells (dark blue) are in the uppermost epidermal stratum and basal keratinocytes (light blue) are in the lowermost epidermal stratum. MC containing cytoplasmic granules, extending microvillar processes, and contacting axons localize between keratinocytes. **(E)** Lateral confocal micrograph of the trunk epidermis in an adult expressing reporters for keratinocytes (*Tg(actb2:LOXP-BFP-LOXP-DsRed)*) and *atoh1a*-expressing cells (*Tg(atoh1a:nls-Eos)*). Dotted boxes indicate areas of magnification in E’ and E’’. **(E’)** Magnification of E showing *atoh1a+* hair cells (HCs) and progenitors within neuromasts (nm) of the posterior lateral line. **(E’’)** Magnification of E showing *atoh1a+* MCs scattered throughout the scale epidermis. **(F)** Lateral and reconstructed cross sectional confocal micrographs of the trunk in an adult expressing reporters for keratinocytes (*Tg(actb2:LOXP-BFP-LOXP-DsRed)*) and *atoh1a*-expressing cells (*Tg(atoh1a:nls-Eos)*) and stained with Alizarin Red S (ARS) to label the mineralized scale matrix. Note that *atoh1a+* MCs localize to the epidermis above scales (arrowhead). **(G)** Lateral confocal micrograph of the scale epidermis in an adult expressing reporters for keratinocytes (*Tg(krt4:DsRed)*) and F-actin within *atoh1a*+ MCs (*Tg(atoh1a:Lifeact-EGFP)*). Note that all *atoh1a+* MCs extend multiple microvilli. **(G’)** Magnification of G with arrowheads indicating individual microvillar processes on the surface of MCs. **(G’’)** Reconstructed cross section along the yellow line in G. MCs localize to the upper epidermal strata as diagrammed in D. Note that *Tg(krt4:DsRed)* (blue) preferentially labels keratinocytes in the upper epidermal strata, but not in the basal cell layer. Scale bars: 1 µm (B,C), 0.1 µm (B’-B’’’), 0.5 µm (C’), 50 µm (E-E’’,F), 10 µm (G) and 5 µm (G’,G’’).

### *atoh1a* reporters label MCs in the adult epidermis

To date, a lack of genetically encoded reagents has hindered in-depth study of anamniote MCs. Since the TEM studies of Whitear and colleagues decades ago, molecular markers have been identified that distinguish mammalian MCs from other epidermal cells. For example, expression of Atoh1 uniquely identifies MCs in rodent skin and is necessary and sufficient for MC development (Morrison et al., 2009; Ostrowski et al., 2015; Van Keymeulen et al., 2009). The zebrafish genome contains three genes (*atoh1a, atoh1b,* and *atoh1c*) encoding Atoh1 homologs (Chaplin et al., 2010; Kani et al., 2010). To determine if the adult epidermis contained cells expressing an Atoh1 homolog, we focused on characterizing the expression pattern of *atoh1a* due to the availability of an enhancer trap line that expresses a nuclear localized version of the photoconvertible fluorescent protein Eos (nls-Eos) from the endogenous *atoh1a* locus (*Tg(atoh1a:nls-Eos)*) (Pickett et al., 2018). Confocal imaging of the adult trunk revealed that *Tg(atoh1a:nls-Eos)* labeled hair cells of the posterior lateral line, which formed tight clusters within neuromasts in interscale regions (Figure 1E,E’). In addition to *atoh1a+* cells of the lateral line, we identified a second, spatially distinct population of *atoh1a+* cells dispersed across the scale surface (Figure 1E,E’’,F). Reconstructed cross-sections showed that this population of *atoh1a+* cells resided within the epidermis above scales (Figure 1F), in a similar axial position to the cells we identified by TEM.

The numerous, actin-rich microvilli that emanate from the MC surface distinguish them from other epidermal cells morphologically (Lane and Whitear, 1977; Toyoshima et al., 1998; Yamashita et al., 1993). To determine whether the dispersed epidermal *atoh1a+* cells extended microvilli, we created an *atoh1a* enhancer trap line that expresses Lifeact-EGFP, a reporter for filamentous actin (Riedl et al., 2008). Similar to *Tg(atoh1a:nls-Eos)*, *Tg(atoh1a:Lifeact-EGFP)* labeled hair cells of the lateral line and inner ear in larvae (Figure 1—figure supplement 1). *atoh1a+* cells were notably absent from regions above the larval eye, yolk sac, or caudal fin (Figure 1—figure supplement 1), where neuroepithelial cells (NECs), a morphologically distinct population of sensory cells, have been described in larval skin (Coccimiglio and Jonz, 2012). Confocal microscopy of the scale epidermis in *Tg(atoh1a:Lifeact-EGFP)* adults revealed actin-rich microvilli densely decorating *atoh1a+* cells in close proximity to neighboring keratinocytes (Figure 1G), further suggesting that the epidermal *atoh1a+* cell population shared key characteristics with the candidate MCs identified by TEM. Immunostaining for Sox2, a transcription factor required for murine MC maturation (Bardot et al., 2013; Perdigoto et al., 2014), demonstrated that the epidermal *atoh1a+* cells expressed Sox2 (Figure 1—figure supplement 2). Together, these results define molecular and cellular properties of a previously uncharacterized epidermal cell population in zebrafish and identify genetic reagents for the study of this cell type. Anticipating the conclusion of our analysis below, we shall hereafter refer to the epidermal *atoh1a+* cells as MCs.

### Somatosensory axons innervate zebrafish MCs, which display neurosecretory and mechanosensory characteristics

We next sought to determine whether zebrafish MCs displayed other key characteristics of MCs defined in mammals, including innervation by somatosensory axons and expression of neurosecretory and mechanosensory machinery.

Our ultrastructural observations suggested that cutaneous axons innervate MCs (Figure 1B). Staining scales with zn-12, a monoclonal antibody that labels several types of peripheral axons (Metcalfe et al., 1990), revealed that >90% of MCs tightly associated with axons (Figure 2A,C). To determine the type of axon(s) innervating MCs, we examined expression of genetically encoded somatosensory axon reporters that we previously characterized in adult scales (Rasmussen et al., 2018). Analysis of reporters for three somatosensory neuron-expressed genes (*p2rx3a*, *p2rx3b,* and *trpa1b*) (Kucenas et al., 2006; Palanca et al., 2013; Pan et al., 2012) demonstrated that somatosensory axons innervated up to 99% of MCs (Figure 2B,C). Consistent with ultrastructural analyses of MCs in the skin of other teleosts (Whitear, 1989), some axons formed ring-like structures that wrapped around MCs and MC-axon contacts frequently contained varicosities or swellings (Figure 2B, inset and 2D-F; Supplemental Video 1). Additionally, we observed examples of axons forming both bouton- and en passant-like contacts with MCs (Figure 2G,H).

**Figure 2.**
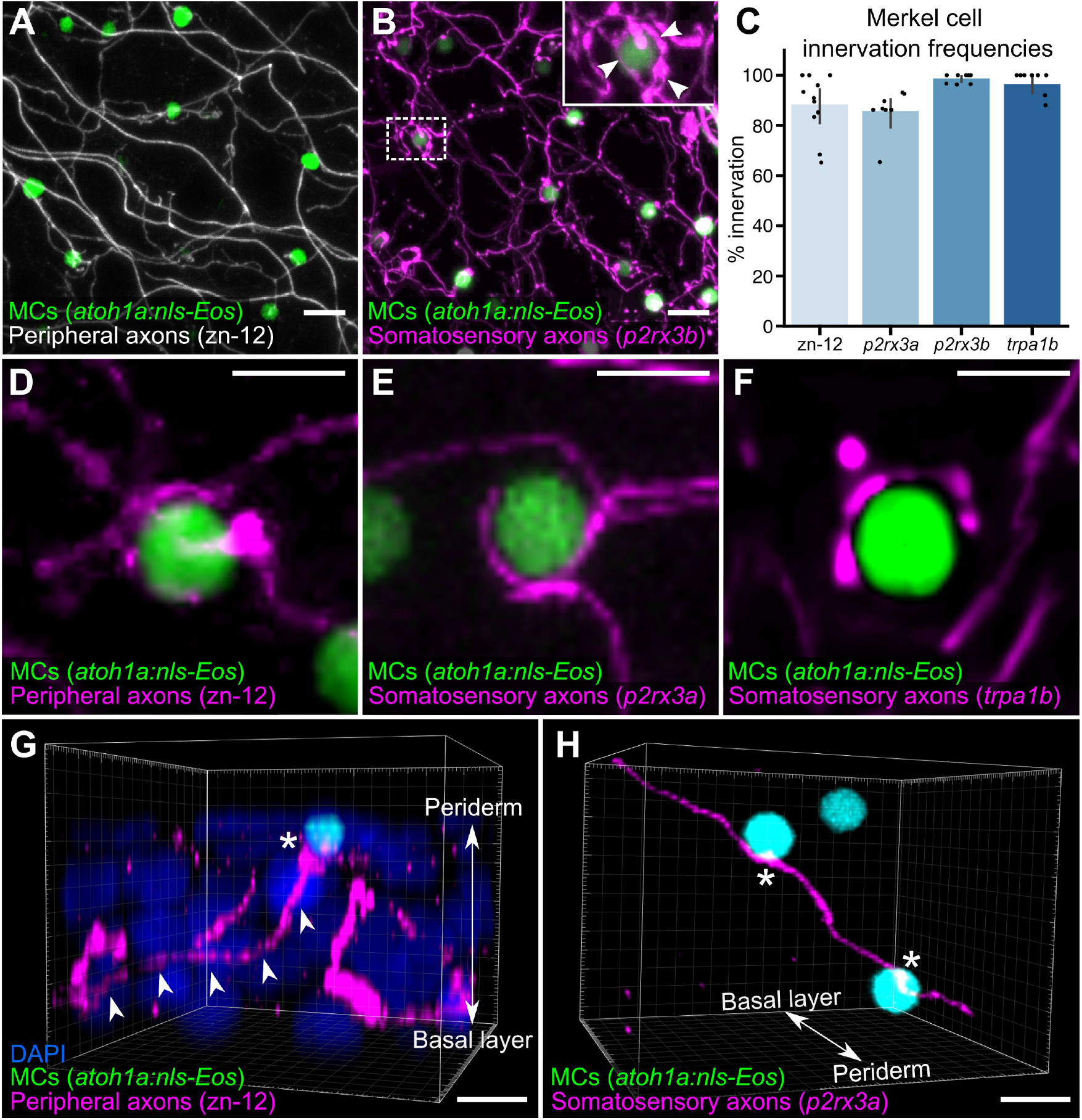
Somatosensory axons innervate MCs in the adult epidermis. **(A)** Lateral confocal micrograph of the scale epidermis from an adult expressing a MC reporter immunostained for peripheral axons (zn-12). **(B)** Lateral confocal micrograph of the scale epidermis showing that somatosensory peripheral axons (*Tg(p2rx3b:EGFP)*) innervate MCs. Inset of dotted region shows axonal varicosities adjacent to a MC (arrowheads). **(C)** Quantification of MC innervation with various axon markers. Each dot represents measurements from an individual scale. Innervation frequencies: zn-12, 91% (284/311 cells; *N*=3 adults); *Tg(p2rx3a>mCherry)*, 86% (196/228 cells; *N*=4 adults); *Tg(p2rx3b:EGFP)*, 99% (225/228 cells; *N*=4 adults); *Tg(trpa1b:EGFP)*, 96% (217/225 cells; *N*=9 adults). **(D-F)** High-magnification confocal micrographs showing examples of somatosensory axons forming extended, ring-like contacts with MCs within the scale epidermis. **(G)** Three-dimensional (3D) reconstruction of an axon (zn-12 immunostaining, arrowheads) forming a bouton-like ending (asterisk) that terminates near a MC. DAPI staining labels epidermal nuclei. **(H)** 3D reconstruction of a single somatosensory axon (*Tg(p2rx3a>mCherry)*) that forms en passant-like contacts (asterisks) with multiple MCs. Scale bars: 10 μm (A, B), 5 μm (D-H).

Based on our observation that MCs contained cytoplasmic granules (Figure 1B), we postulated that they would display neurosecretory characteristics. We began by staining scales with an antibody against synaptic vesicle glycoprotein 2 (SV2), a component of secretory vesicle membranes (Buckley and Kelly, 1985). Essentially all MCs contained SV2-positive structures (Figure 3A), suggesting they express neurosecretory machinery that may contain neurotransmitter(s). Indeed, immunostaining revealed that MCs expressed serotonin (5- hydroxytryptamine; 5-HT) (Figure 3B), similar to mammalian MCs (Chang et al., 2016; English et al., 1992; García-Caballero et al., 1989). Both 5-HT and SV2 appeared in a speckled pattern within MCs (Figure 3A,B), consistent with a vesicular localization.

**Figure 3.**
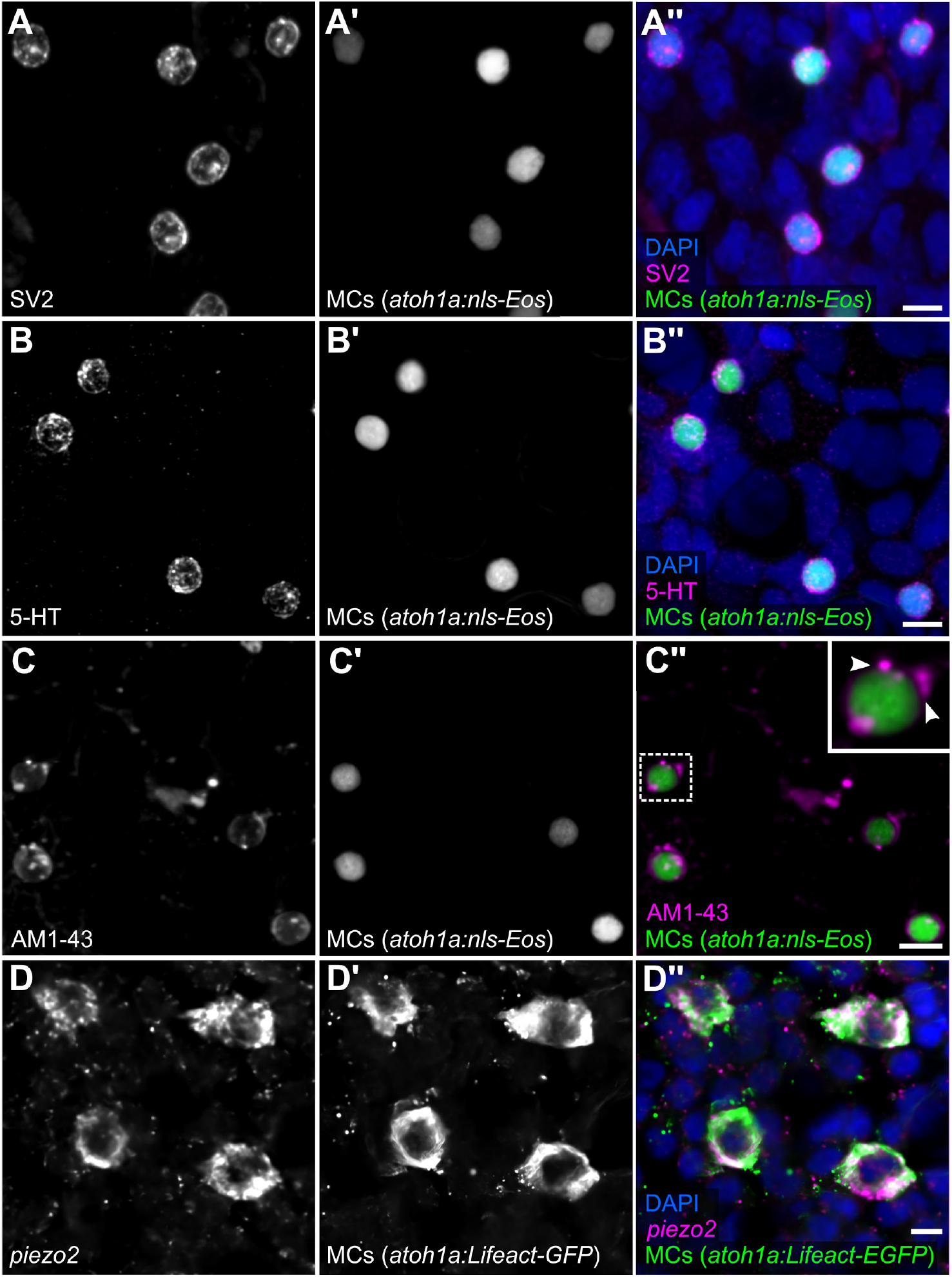
MCs in the adult epidermis express neurosecretory and mechanosensory machinery. **(A,B)** Anti-SV2 (A-A’’) or anti-5-HT (B-B’’) immunostaining of the scale epidermis from an adult expressing a MC reporter. Note the punctate localization of SV2 and 5-HT staining in MCs, consistent with a vesicular localization. 96% of MCs (178/185) were SV2+. 98% of MCs (326/332) were 5-HT+. Cells analyzed from *n*=3 scales from *N*=2 adults (25-27 mm SL). DAPI labels epidermal nuclei. **(C)** Scale epidermis from an adult expressing a MC reporter stained with AM1-43. 98% of MCs (90/92) were AM1-43+. Cells analyzed from *n*=6 scales from *N*=2 adults. Inset of dotted region shows puncta within a MC labeled by AM1-43 (arrowheads). (**D)** Fluorescent in situ hybridization with an antisense probe against *piezo2.* 99% of MCs (246/248) were *piezo2*+. Cells analyzed from *n*=3 individual scales from *N*=2 adults (23-27 mm SL). Scale bars: 5 µm.

Do zebrafish MCs exhibit properties consistent with mechanosensory function? To address this question, we began by staining scales with AM1-43, an activity-dependent fluorescent styryl dye that labels a variety of sensory cells, including mammalian MCs (Meyers et al., 2003). Following a short preincubation, AM1-43 robustly stained MC membranes and punctate structures reminiscent of vesicular compartments (Figure 3C), suggestive of ion channel expression in MCs (Meyers et al., 2003). Mammalian MCs express the mechanically activated cation channel Piezo2, which is required for MC mechanosensory responses (Ikeda et al., 2014; Maksimovic et al., 2014; Woo et al., 2014). Fluorescent in situ hybridization with an antisense probe against *piezo2* strongly labeled MCs in adult scales (Figure 3D). Together, these data suggest that somatosensory peripheral axons innervate adult MCs, which possess neurosecretory and mechanosensory properties.

### MCs arise from basal keratinocyte precursors

What are the precursors of MCs in zebrafish? Analysis of MC progenitors have come to conflicting results in avians and rodents: quail/chick chimeras suggest a neural crest origin for avian MC (Grim and Halata, 2000), whereas Cre-based lineage tracing studies in mouse demonstrate an epidermal origin (Morrison et al., 2009; Van Keymeulen et al., 2009).

To investigate a possible neural crest origin, we crossed a Cre driver expressed in neural crest progenitors (*Tg(sox10:Cre)*; (Kague et al., 2012)) to a reporter transgene that stably expresses DsRed upon Cre-mediated recombination from a quasi-ubiquitous promoter (*Tg(actb2:LOXP-BFP-LOXP-DsRed)*; (Kobayashi et al., 2014)) (Figure 4A). DsRed+ neural crest-derived cell types, such as Schwann cells, appeared along scales, indicative of successful recombination (Figure 4B). However, we observed <0.5% colocalization between the neural crest lineage trace and a MC reporter (Figure 4B’,E). Based on these results, we concluded that zebrafish MCs likely derive from a non-neural crest lineage.

**Figure 4.**
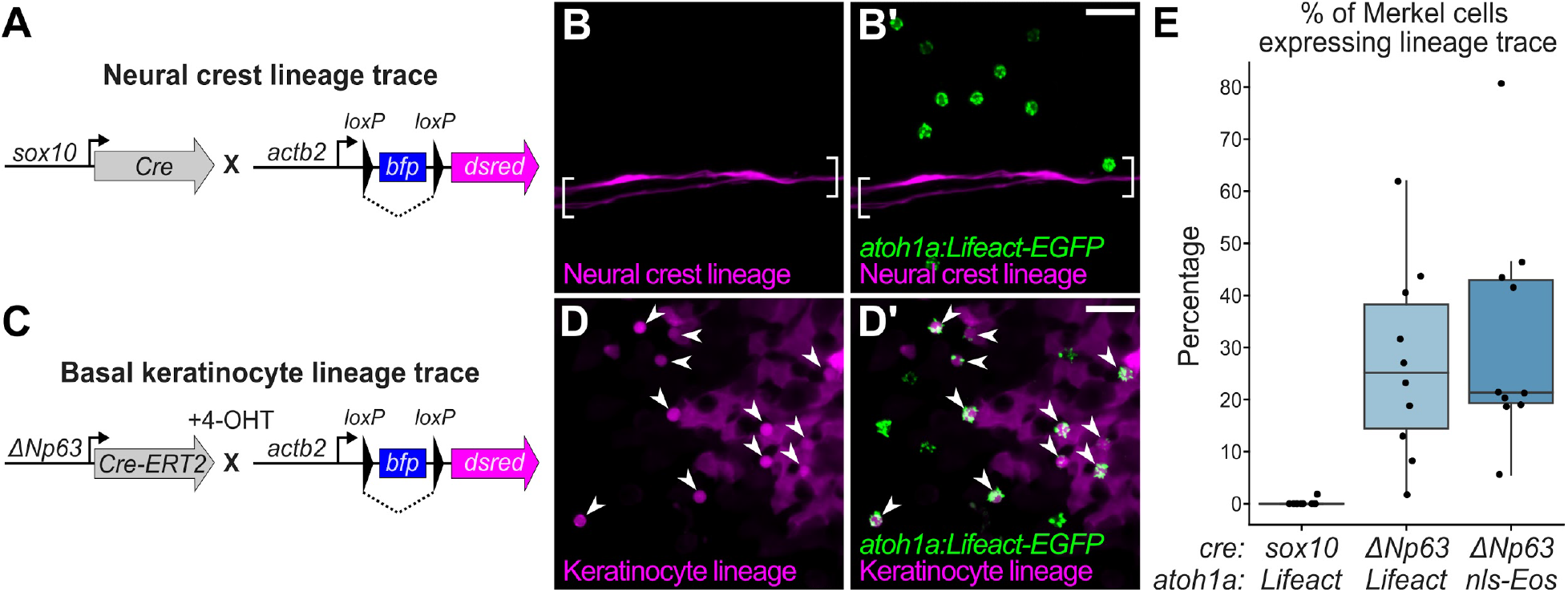
Merkel cells derive from the basal keratinocyte lineage. **(A)** Schematic of Cre-based neural crest lineage tracing strategy. **(B)** Confocal micrograph of the scale epidermis in an adult expressing neural crest lineage (*Tg(sox10:Cre); Tg(actb2:LOXP-BFP-LOXP-DsRed)*) and MC (*Tg(atoh1a:Lifeact-EGFP)*) reporters. Brackets denote Schwann cells associated with a nerve along a scale radius. **(C)** Schematic of Cre-based basal keratinocyte lineage tracing strategy. **(D)** Confocal micrograph of the scale epidermis in an adult expressing basal keratinocyte lineage (*TgBAC(ΔNp63:Cre-ERT2); Tg(actb2:LOXP-BFP-LOXP-DsRed)*) and MC (*Tg(atoh1a:Lifeact-EGFP)*) reporters, which was treated with 4-OHT at 1 dpf. Arrowheads indicate MCs labeled by the basal keratinocyte lineage reporter. Note that recombination is not complete, therefore not all MCs express the lineage reporter. **(E)** Boxplots of the percentage of MCs expressing the lineage tracing reporters diagrammed in panels A and C. Each dot represents an individual scale. Overall percentage of MCs expressing lineage trace reporters: *sox10/Lifeact*, 0.3% (1/323 cells; *N*=6 adults); *ΔNp63/Lifeact*, 29.7% (299/1005 cells; *N*=6 adults); *ΔNp63/nls-Eos*, 32.3% (386/1195 cells; *N*=4 adults). Scale bars: 20 µm.

To investigate a possible epidermal origin, we considered basal keratinocytes, an epidermal-resident stem cell population, the most likely candidate progenitors. To follow this lineage, we engineered a transgene to express a tamoxifen-inducible Cre recombinase from regulatory sequences of *ΔNp63* (*TgBAC(ΔNp63:Cre-ERT2)*), a basal keratinocyte marker (Bakkers et al., 2002; Lee and Kimelman, 2002). We crossed this transgene to the Cre reporter transgene and treated embryos with 4-OHT at 1 day post-fertilization (dpf) to induce Cre-ERT2 activity, which resulted in permanent DsRed expression in basal keratinocytes and their derivatives (Figure 4C,D; Figure 4—figure supplement 1). 4-OHT-treated animals showed extensive co-labeling between the basal keratinocyte lineage trace and a MC reporter (Figure 4D’,E). These observations strongly support a basal keratinocyte origin of zebrafish MCs.

### MCs continuously turn over in adult skin

The longevity and turnover of murine MCs is controversial. Several studies concluded that MC numbers fluctuate with hair cycle stages (Marshall et al., 2016; Moll et al., 1996; Nakafusa et al., 2006), while another found no correlation (Wright et al., 2017). To determine the turnover rate of zebrafish MCs, we photoconverted small regions of the scale epidermis in *Tg(atoh1a:nls-Eos)* adults and tracked individual cells over time. Exposure to UV light irreversibly photoconverts nls-Eos, allowing us to distinguish pre-existing cells (containing photoconverted nls-Eos) from newly added cells (without photoconverted nls-Eos) (Figure 5A-C). By longitudinally tracking individual fish over the course of 28 days, we found a decrease of ∼15% of the photoconverted MCs every 7 days (Figure 5D). In addition to the gradual loss of MCs over time, we noted a steady addition of new MCs, resulting in a nearly constant total cell number (Figure 5E). Thus, MCs undergo constant cell loss and renewal in adult skin, albeit at a slower rate than *atoh1a*-expressing hair cells of the lateral line (Cruz et al., 2015).

**Figure 5.**
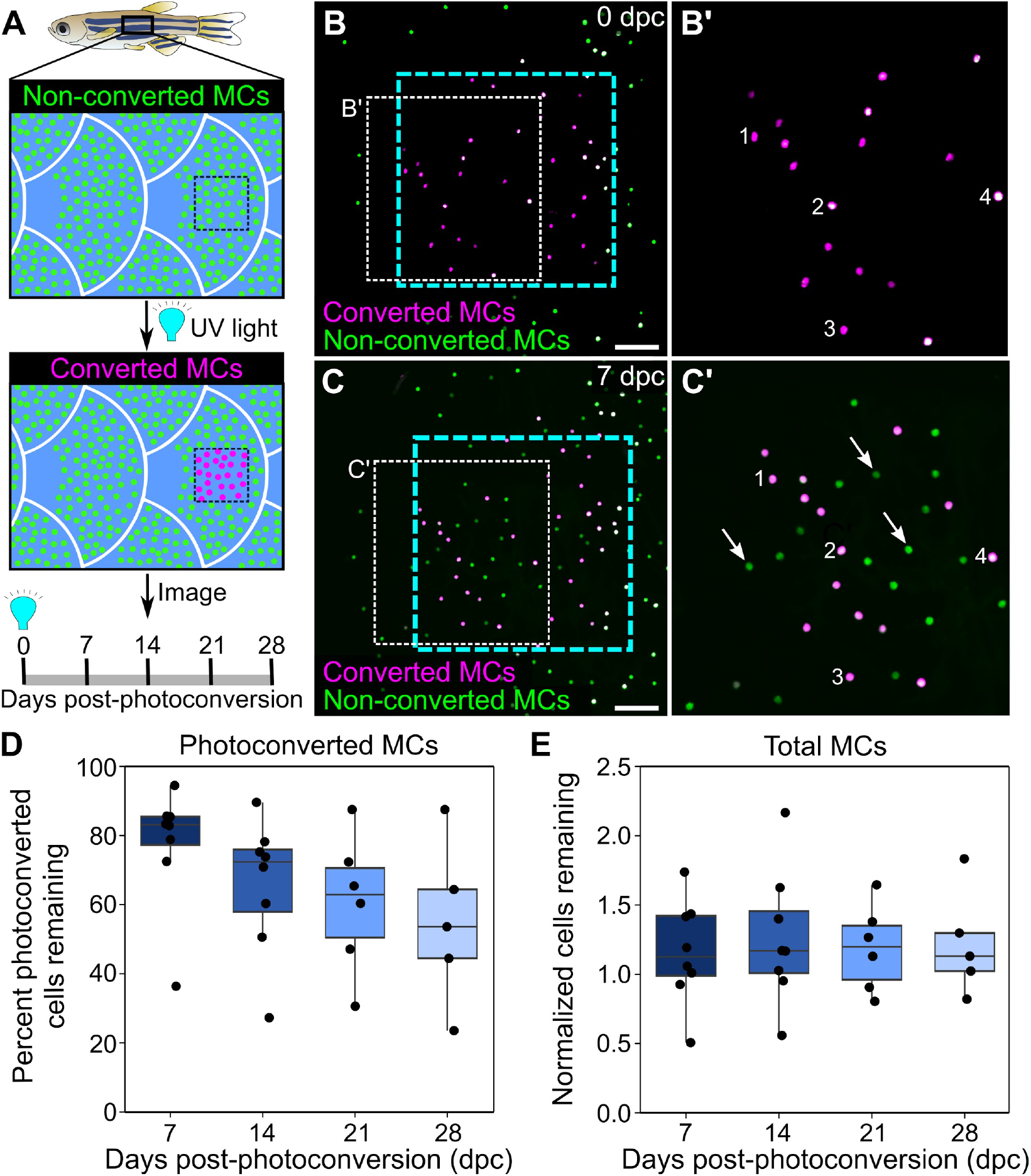
Homeostatic replacement of MCs in the adult epidermis. **(A)** Illustration of the photoconversion experiment showing the epidermis (blue), non-converted MCs (green), and converted MCs (magenta) after exposure of a region of the scale epidermis to UV light. **(B,C)** Representative images of MCs labeled by *Tg(atoh1a:nls-Eos)* at 0 (B) or 7 (C) days post-conversion (dpc) from a single adult. Cyan dotted box indicates the photoconverted region. White dotted box indicates the area magnified in B’, C’. **(B’,C’)** Numbers label examples of individual cells present at 0 and 7 dpc. Arrows indicate examples of newly added cells, which appear green due to the presence of non-converted nls-Eos (green) and absence of converted nls-Eos (magenta). **(D)** Boxplots of the percent of photoconverted MCs remaining compared to 0 dpc. Each dot represents an individual fish. **(E)** Boxplots of the total number of MCs (converted + non-converted) present at each day compared to 0 dpc. Each dot represents an individual fish. Scale bars: 50 μm.

### MCs are widely distributed across the body, in compartment-specific patterns

MCs localize to specific regions of mammalian skin, such as in crescent-shaped touch domes adjacent to hair follicles in hairy skin and at the bottom of rete ridges in glabrous skin (Boot et al., 1992; Fradette et al., 1995; Iggo and Muir, 1969; Lacour et al., 1991). To determine the distribution pattern of zebrafish MCs, we used confocal microscopy to survey multiple regions of the adult skin. In addition to the MCs found on the trunk, MCs appeared in the epidermis above the eyes, gill covers (opercula), and fins (Figure 6A-E). While MC morphology was similar across the skin compartments (Figure 6A-E, insets), MC densities and spatial distributions varied across skin compartments (Figure 6F). For example, MCs were distributed uniformly across the eye (Figure 6B). By contrast, in the caudal fin, MCs localized specifically to the epidermis above bony rays and in medial regions of the interray epidermis between bony rays (Figure 6E). Along the trunk, MCs appeared in patches, similar to the pattern of dermal scales beneath the epidermis (Figure 6D). Altogether, our results demonstrate that MCs are widely distributed across the adult zebrafish skin and localize in specific patterns in each skin compartment.

**Figure 6.**
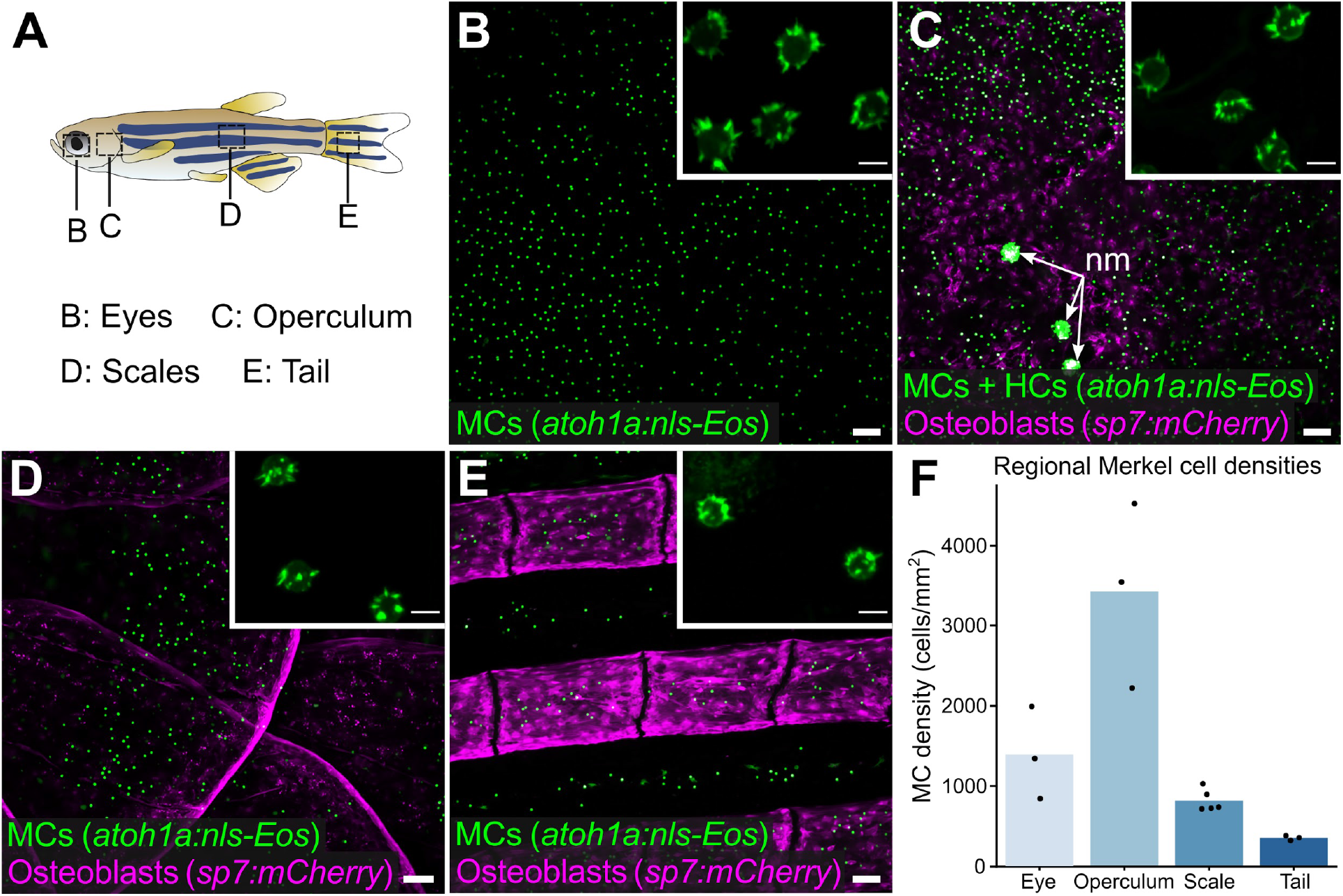
MCs are widely distributed across the skin, in compartment-specific patterns. **(A)** Illustration indicating the regions imaged in adult zebrafish. **(B-E)** Lateral confocal micrographs of MCs in the regions from animals expressing the indicated reporters. The regions imaged are indicated in A. Insets show MCs expressing *Tg(atoh1a:Lifeact-EGFP)* have a similar morphology across skin compartments. nm, neuromasts of the posterior lateral line. **(F)** Quantification of MC densities in the specified regions. Each dot represents an individual fish (27-29 mm SL). Scale bars: 50 μm (B-E), 5 μm (B-E, insets).

### Trunk MCs develop concomitant with dermal appendage morphogenesis

To examine the mechanisms that generate a compartment-specific MC pattern, we focused on the trunk skin because of its molecular and cellular similarities to murine hairy skin (Aman et al., 2018; Harris et al., 2008). Both during ontogeny and at post-embryonic stages, murine MCs associate with primary (guard) hairs, a subclass of dermal appendages (Jenkins et al., 2019; Nguyen et al., 2018). Based on these studies in mice, and our previous work showing that epidermal diversification and somatosensory remodeling coincides with scale development in zebrafish (Rasmussen et al., 2018), we postulated that MCs would appear during squamation (scale formation).

Zebrafish post-larval development is staged by standard length (SL) in millimeters (mm) (Parichy et al., 2009). Squamation begins at ∼9 mm SL (Figure 7A) (Aman et al., 2018; Harris et al., 2008; Sire et al., 1997a). Using reporters that label MCs and scale-forming osteoblasts, we observed only rare MCs in the epidermis prior to the onset of squamation (Figure 7B). By contrast, MC density rapidly increased between 10-15 mm SL, a period of active scale growth (Figure 7C-F). The density and number of MCs positively correlated with scale area (Figure 7G,H). These data indicate that MC development coincides with dermal appendage growth along the trunk.

**Figure 7.**
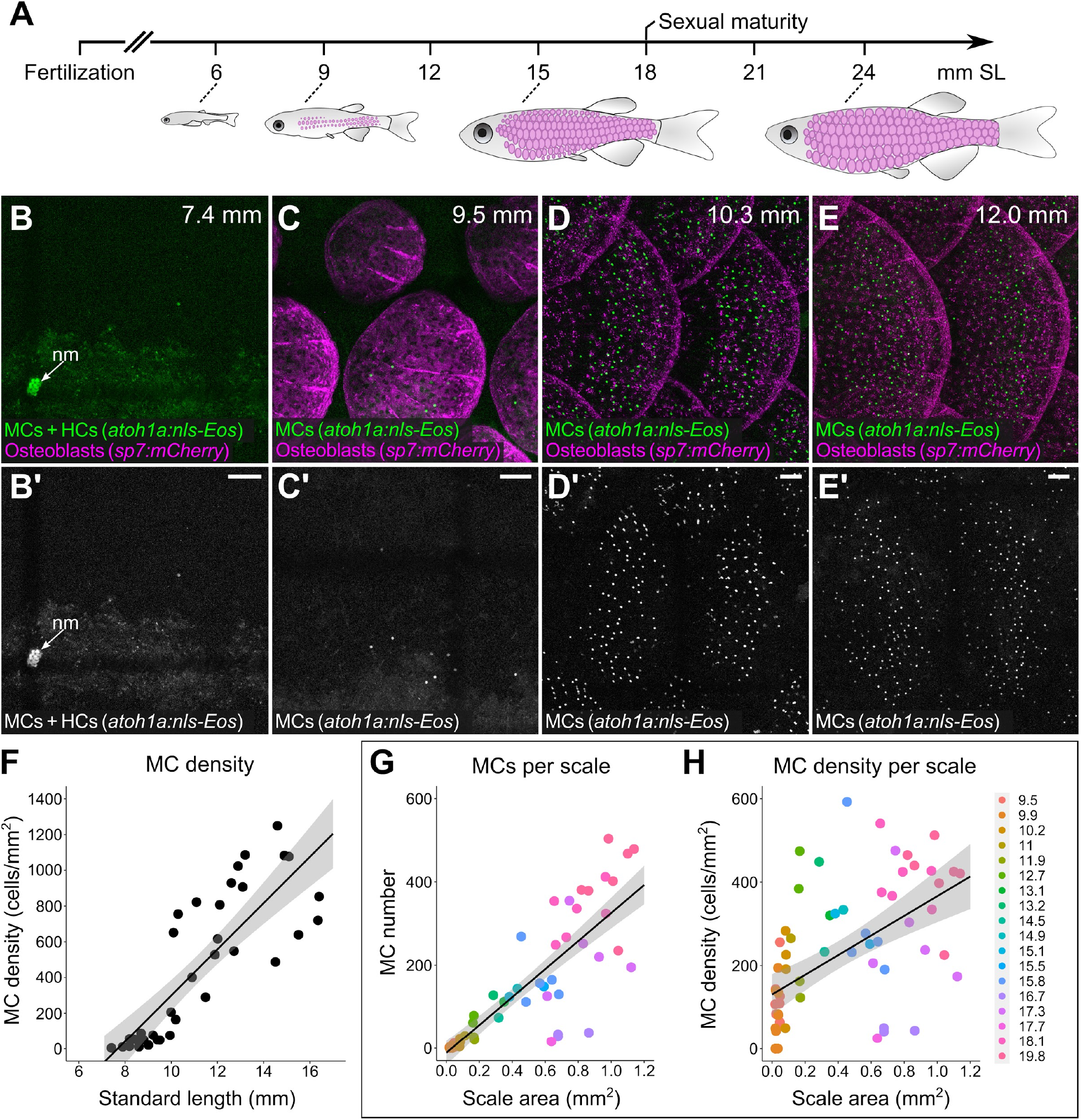
MCs develop concomitant with dermal appendage morphogenesis. **(A)** Abbreviated zebrafish developmental timeline showing standard length (SL) in millimeters. Developing scales are drawn in magenta below the approximate corresponding stage. **(B-E)** Representative lateral confocal micrographs of MCs and osteoblasts along the trunk at the indicated stages. Note that MCs increase in number and density as scale-forming osteoblasts develop below the epidermis. nm, neuromast of the posterior lateral line. **(F)** Quantification of MC density according to SL. Each dot represents an individual fish. **(G,H)** Quantification of the number (G) or density (H) of MCs relative to scale area. Each dot represents an individual scale. Dot colors represent animal SL as indicated in the legend. Shading indicates a 95% confidence interval around the linear regression lines in F-H. Correlation coefficients (R^2^): 0.33 (F), 0.73 (G), 0.31 (H). Scale bars: 50 μm.

### Ectodysplasin signaling promotes trunk MC development

Since appearance of MCs in the trunk epidermis tightly correlated with scale growth, we examined the consequences of blocking dermal appendage morphogenesis on MC development. Ectodysplasin (Eda) signaling regulates the formation of many types of skin appendages, including mammalian hair follicles and zebrafish scales (Biggs and Mikkola, 2014; Harris et al., 2008). To determine whether MC development requires Eda-dependent signals, we measured MC density in animals homozygous for a presumptive null allele of *eda* that do not develop scales (*eda^dt1261/dt1261^*; hereafter *eda^−/−^*) (Harris et al., 2008). Immediately prior to squamation, we found that there was no difference in MC density between *eda* mutants and sibling controls (Figure 8A,B,G). However, after the onset of squamation, *eda* mutants had significantly fewer MCs, a difference that persisted into adulthood (Figure 8A-G). In addition to the decrease in cell density, we observed a dramatic change in the spatial distribution of MCs across the epidermis in *eda* mutants compared to controls (Figure 8H). Specifically, in siblings, MCs appeared in patches corresponding to the location of the underlying scales (Figure 8C,H). By contrast, the few MCs that developed in *eda* mutants were distributed uniformly across the trunk (Figure 8D,H). Although we found a decrease in MC density in the trunk skin of the mutants, we observed no change in MC density above the eye or operculum (Figure 8I), suggesting that the reduced density was specific to the trunk skin.

**Figure 8.**
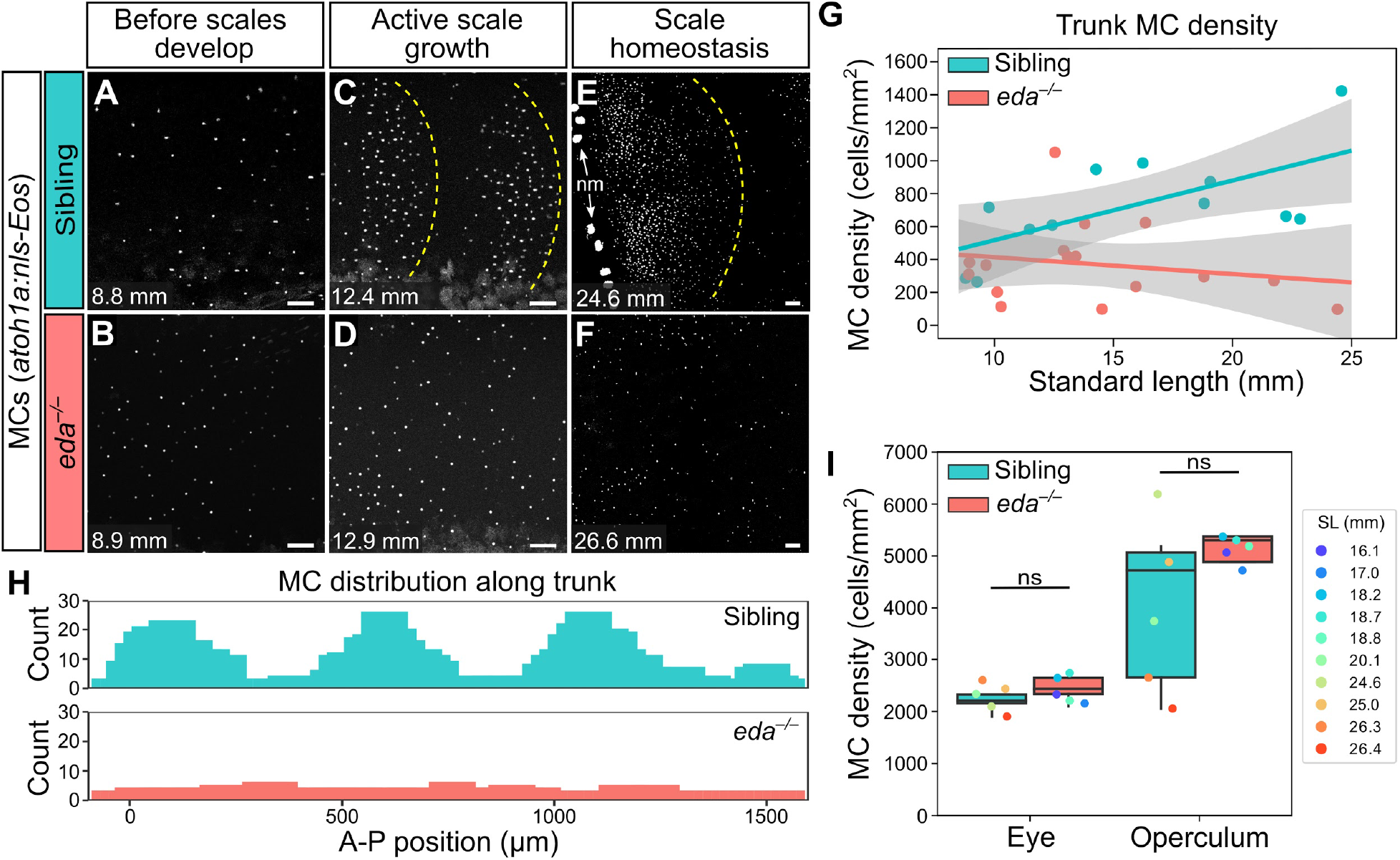
Preventing dermal appendage development decreases MC density in trunk, but not facial, skin. **(A-F)** Representative confocal images of MCs in juveniles of the indicated genotypes at the indicated stages. Dotted yellow lines indicate posterior scale boundaries. nm, neuromasts of the posterior lateral line. **(G)** Quantification of MC density in the trunk skin relative to standard length. Grey shading indicates a 95% confidence interval around the linear regression lines. The difference between genotypes was significant above 12.5 mm SL (*P* < 0.05, Johnson-Neyman Technique). Each dot represents an individual animal. **(H)** Histograms of the distribution of trunk MCs along a rectangular segment encompassing 3 scales in a sibling and an identically sized region in an *eda* mutant (18-19 mm SL). **(I)** Boxplots of MC densities in the epidermis above the eye or operculum in animals of the indicated genotypes. ns, not significant (eye, *P*=0.21; operculum, *P*=0.14; Mann-Whitney test). Scale bars: 50 μm (A-F).

The decreased MC density in *eda* mutant trunk skin could be due to decreased cell addition, increased cell turnover rate, or a combination of the two. Using *in vivo* photoconversion, we found that the rate of MC addition was significantly reduced in *eda* mutants compared to siblings (Figure 8—figure supplement 1A-D). Additionally, the rate of cell loss was higher in mutants compared to siblings (Figure 8—figure supplement 1E). Thus, our observations indicate that the decrease in MC cell density in *eda* mutants is likely due to both reduced MC production and increased MC turnover. Together, these data suggest that Eda signaling is required for MC development, maintenance, and distribution along the trunk.

### Altering dermal appendage shape and size redistributes MCs

Since blocking dermal appendage formation inhibited MC development, we next examined the consequences of altering dermal appendage size and shape on MC patterning. Zebrafish scale morphogenesis is regulated by Fibroblast growth factor (FGF) signaling (Aman et al., 2018; Daane et al., 2016; De Simone et al., 2021; Rohner et al., 2009). To determine whether alterations to scale patterning impacted MC development, we examined animals heterozygous for an allele of *hagoromo (hag; fgf8a^dhiD1Tg/+^)*, which results in *fgf8a* overexpression in the post-embryonic skin due to a viral insertion near the *fgf8a* locus (Amsterdam et al., 2009). An independent allele of *hag (fgf8a^dhi4000Tg/+^)* was previously shown to result in large, disorganized sheets of scale-forming osteoblasts during squamation (Aman et al., 2018). *fgf8a^dhiD1Tg/+^* juveniles showed dramatic variability in scale size and shape, with both smaller and larger scales compared to the remarkably uniformly patterned scales observed in sibling controls (Figure 9A-D; Figure 9—figure supplement 1A-C). We found no significant differences in MC density between the genotypes (Figure 9— figure supplement 1D,E). Nevertheless, the distribution of MCs tracked with the altered scale size and shape in the mutants, suggesting the MC pattern is not predetermined (Figure 9). Based on these data, we concluded that altering dermal appendage morphogenesis is sufficient to redistribute MCs within the trunk skin compartment.

**Figure 9.**
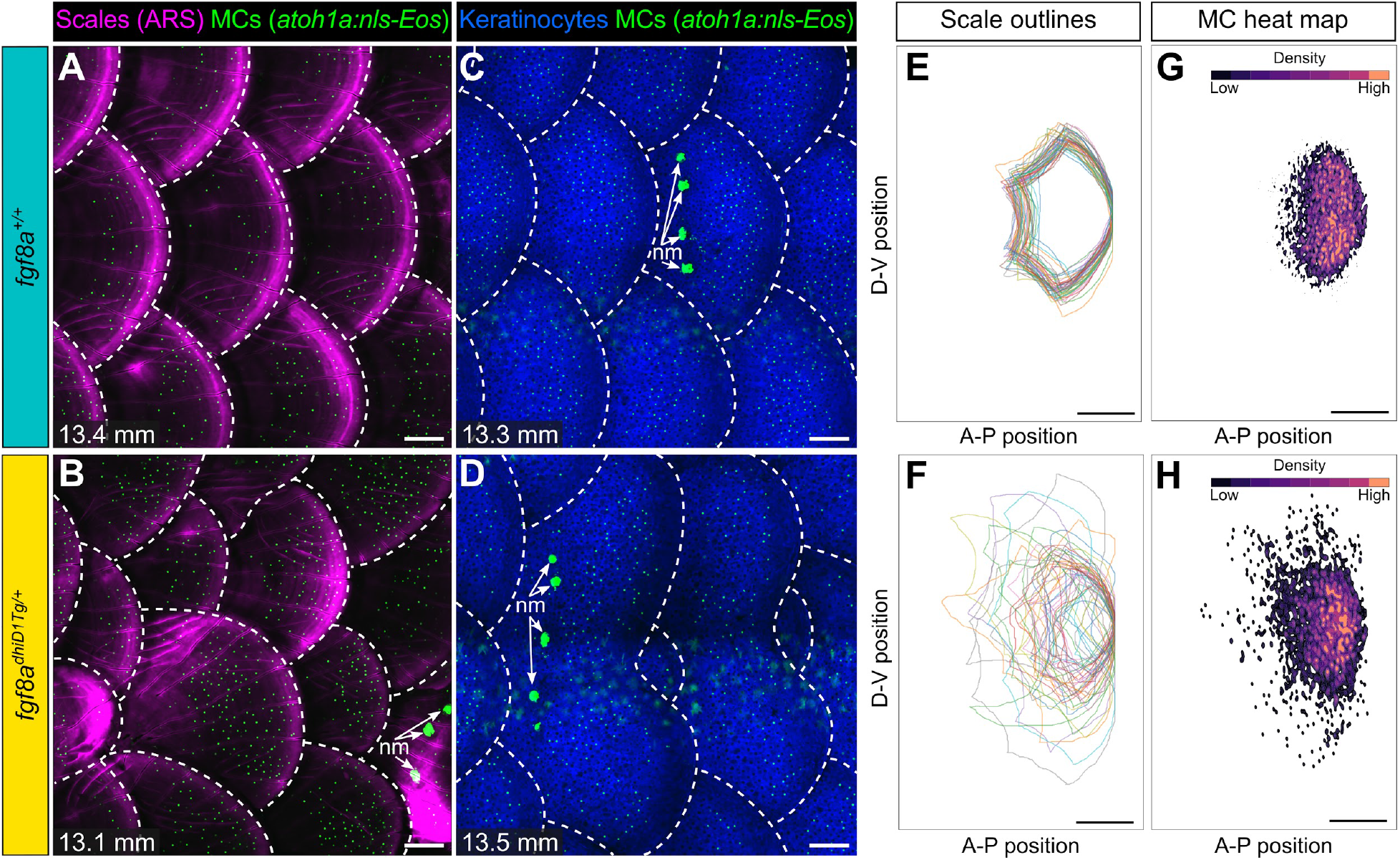
Altering dermal appendage patterning redistributes MCs. (**A-D)** Representative images of juvenile of the indicated genotypes expressing a MC reporter and stained with ARS to visualize scales (A,B) or co-expressing MC and keratinocyte (*Tg(krt4:DsRed)*) reporters (C,D). Dotted lines indicate scale boundaries. nm, neuromasts of the posterior lateral line. **(E-H)** Tracings of scale outlines (E,F) and density plots of MC position (G,H) from juvenile animals (11.6-14.7 mm SL) of the indicated genotypes. Scales tracings were aligned at the dorsal-ventral midpoint of the posterior scale margin. Note the variability in scale shape and size and corresponding increased spread of MC position in *fgf8a^dhiD1Tg/+^*juveniles compared to sibling controls. Scale bars: 100 μm (A-D), 200 μm (E-H).

## DISCUSSION

Here, we discover a zebrafish epidermal cell type that we classify as a MC based on ultrastructural criteria (Whitear, 1989). We further present several lines of evidence that suggest zebrafish MCs share molecular, cellular, and lineage properties with mammalian MCs. First, we show that zebrafish MCs express the transcription factors Atoh1a and Sox2, the orthologs of which uniquely mark MCs in mammalian skin (Maricich et al., 2009; Nguyen et al., 2018; Ostrowski et al., 2015; Van Keymeulen et al., 2009). Second, zebrafish MCs extend numerous short, actin-rich microvilli and complex with somatosensory axons, classic morphological hallmarks of MCs (Mihara et al., 1979; Smith, Jr, 1977; Toyoshima et al., 1998). Third, Cre-based lineage tracing revealed that basal keratinocytes give rise to zebrafish MCs, akin to studies in mouse (Morrison et al., 2009; Van Keymeulen et al., 2009). Fourth, we demonstrate that zebrafish MCs contain neurosecretory machinery and express the neurotransmitter serotonin, release of which has been proposed to regulate somatosensory responses to touch (Chang et al., 2016; Chang and Gu, 2020; English et al., 1992). Finally, we show that zebrafish MCs express the cation channel Piezo2, which is cell-autonomously required for MC mechanosensory function (Ikeda et al., 2014; Maksimovic et al., 2014; Woo et al., 2014). Importantly, our results extend previous histological studies of MCs in various teleost fish (Hartschuh and Weihe, 1980; Lane and Whitear, 1977; Whitear, 1989; Whitear and Lane, 1981; Zachar and Jonz, 2012) by identifying the first genetically encoded reagents for the study of this cell type in zebrafish.

### Teleost Merkel cells and somatosensory physiology

In mammalian skin, the MC-neurite complex regulates slowly adapting type I responses to light touch (Iggo and Muir, 1969; Ikeda et al., 2014; Maksimovic et al., 2014; Maricich et al., 2009; Woo et al., 2014). Although physiological studies of somatosensory responses in adult zebrafish have not been reported, extracellular recordings in adult rainbow trout demonstrated that a subset of somatosensory neurons exhibited slowly adapting responses to mechanical skin stimulation (Ashley et al., 2007, 2006; Sneddon, 2003). We postulate that the slowly adapting responses to mechanical skin stimulation in adults requires MCs. Nevertheless, the exact physiological roles of teleost MCs in regulating somatosensory responses and resulting behaviors remain unknown and will require the development of tools to selectively ablate and activate MCs. Interestingly, recordings from zebrafish Rohon-Beard neurons, a transient larval somatosensory population, suggest they have rapidly, but not slowly, adapting mechanosensory responses (Katz et al., 2021). Together these studies correlate with our finding that MCs develop at post-larval stages and suggest that the teleost somatosensory system undergoes significant functional maturation during the juvenile period.

What are the subtypes of somatosensory neurons in fish and how do they correspond to MC innervation? Several studies have identified molecularly distinct subsets of somatosensory neurons through mRNA, protein, and transgene expression analysis in larvae (Faucherre et al., 2013; Gau et al., 2017, 2013; Kucenas et al., 2006; Palanca et al., 2013; Pan et al., 2012; Patten et al., 2007; Slatter et al., 2005). Adult trout somatosensory neurons have been classified based on their responses to mechanical, chemical, and thermal stimuli (Ashley et al., 2007, 2006; Sneddon, 2003). However, to date, a detailed molecular characterization of the diversity of somatosensory subtypes present in adult fish has not been performed. Our data suggest that somatosensory neurons expressing reporters for *p2rx3a, p2rx3b*, or *trpa1b* innervate MCs. Whether these neurons represent a dedicated class of MC-innervating neurons remains unknown. The development of Cre drivers for specific somatosensory subtypes (Bai et al., 2015; Li et al., 2011; Luo et al., 2009; Rutlin et al., 2014; Zylka et al., 2005) and single-cell transcriptional profiling (Sharma et al., 2020; Usoskin et al., 2015; Zeisel et al., 2018) have been fruitful in characterizing the diversity of somatosensory neurons in mammals. The application of these technologies to the teleost somatosensory system is an interesting avenue for further investigation.

### Merkel cell lineage and homeostasis

The developmental lineage of MCs has been a long-standing question with both epidermal and neural crest origins posited (Hartschuh et al., 1986). Our Cre-based lineage tracing identified basal keratinocytes as MC progenitors. These results extend previous studies in the zebrafish epidermis showing that basal keratinocytes serve as precursors for diverse post-larval cell types, including periderm (superficial epidermis) and immune cells (Lee et al., 2014; Lin et al., 2019). Although previous work in mouse unambiguously identified *keratin 14*-expressing basal keratinocytes as MC precursors (Morrison et al., 2009; Van Keymeulen et al., 2009), the precise nature of murine MC progenitors varies across skin compartments (Nguyen et al., 2019). Future studies characterizing the molecular properties and cellular behaviors of zebrafish MC precursors will be informative for identifying conserved properties of skin stem cells.

The turnover of MCs in mammalian skin has been a source of controversy. Several studies reported that MC numbers fluctuate with the natural hair cycle in mouse (Marshall et al., 2016; Moll et al., 1996; Nakafusa et al., 2006). By contrast, Wright et al. (2017) found no evidence for changes in MC density based on stages of the hair cycle and demonstrated that MCs could live for months. These types of analyses in murine skin have relied either on histology, which limits tissue sampling, or required use of advanced (2-photon) microscopy in combination with hair shaving, a mild form of skin injury. Using photoconversion and confocal imaging, we non-invasively tracked individual MCs during normal skin homeostasis *in vivo* for weeks. We found that trunk MCs have a steady turnover in adult animals, with a half-life of approximately 1 month. Additionally, this further distinguishes MCs from hair cells in the adult lateral line, which have a shorter half-life (Cruz et al., 2015). Whether MC turnover varies at different stages of development, across skin compartments, or following skin insults will require further study.

### Merkel cell distribution and patterning

Regionally specific sensory structures allow our skin to distinguish tactile inputs with remarkable acuity (Corniani and Saal, 2020). For example, MC densities vary greatly across human skin compartments, with the highest numbers found in particularly sensitive regions such as fingertips and lips (Boot et al., 1992; Lacour et al., 1991). We observed that MCs populate several major skin compartments and have regional-specific densities in adult zebrafish, with the highest densities found in the face (above the eye and operculum). We speculate this may bestow the juvenile and adult skin with the ability to detect innocuous tactile inputs across almost the entire body surface, with perhaps the greatest sensitivity along facial structures.

Although most studies of MC development have centered on the formation of MC aggregates in the touch domes of murine hairy skin, MCs are found in a range of distribution patterns in other types of skin. For example, MCs are found as dispersed, single cells arrayed across the skin of human toe pads (Boot et al., 1992). Similarly, we found that MCs have a dispersed, rather than clustered, pattern in all skin compartments examined. Few studies have addressed how MCs adopt specific distributions, and zebrafish present a promising model to understand mechanisms of MC pattern formation *in vivo*.

### Dermal appendages and Merkel cell development

Our developmental analysis showed that trunk MC density rapidly increases during dermal appendage morphogenesis. Previous genetic analysis in mouse hairy skin revealed that MC development requires Eda signaling (Vielkind et al., 1995; Xiao et al., 2016). We show that zebrafish *eda* mutants have decreased MC density in trunk, but not facial, skin. These observations suggest that MC development in mouse and zebrafish likely share similar genetic pathways, akin to the shared molecular and cellular mechanisms that regulate dermal appendage formation (Aman et al., 2018; Biggs and Mikkola, 2014; Daane et al., 2016; Harris et al., 2008; Rohner et al., 2009). They further support a model whereby MC development requires compartment-specific signals, akin to recent observations on MC development in mouse hairy and glabrous skin (Nguyen et al., 2019). By taking advantage of the ability to image large skin areas in intact zebrafish, we show that *eda* mutants have altered MC distribution compared to controls. Furthermore, we use *in vivo* photoconversion to demonstrate the reduction in MC density is largely due to decreased production, but also reflects increased turnover. One explanation for these results is that Eda signaling regulates the differentiation of MC progenitors. Alternatively, since *eda* mutants have decreased epidermal innervation (Rasmussen et al., 2018), MC development may require somatosensory neuron-derived signals.

We found that a gain-of-function allele of *fgf8a* leads to a change in the overall size and shape of scales. Intriguingly, the MC distribution modifies to accommodate the altered scale size and shape in the *fgf8a* mutants but retains the same MC density as sibling controls. This result suggests that the number of MCs per scale is not predetermined, but rather is titrated relative to appendage size. How are MCs able to populate the much larger scales in *fgf8a* mutants? Does the size of the MC progenitor domain expand with increases in scale size? Are MCs, or their progenitors, able to migrate to their final destination? Distinguishing between these possibilities will require tracking the behaviors of MCs and their progenitors *in vivo*.

### Summary

Our results establish a promising new system to investigate MC biology. This model will allow for the identification of deeply conserved mechanisms used to regulate vertebrate MC biology. Furthermore, the advantages of zebrafish—such as non-invasive *in vivo* imaging, genetic and chemical screens, and high regenerative capacity—will complement the strengths of rodent models. Specifically, the ability to track individual cells over time has the potential to answer key and long-standing questions surrounding MC biology, including how the MC-neurite relationship is established, how MCs interact with neighboring cell types, and their progenitor dynamics. Addressing these questions, as well as potential novel insights provided by the zebrafish system, represent exciting directions for future research.

## MATERIALS AND METHODS

### Key Resource Table

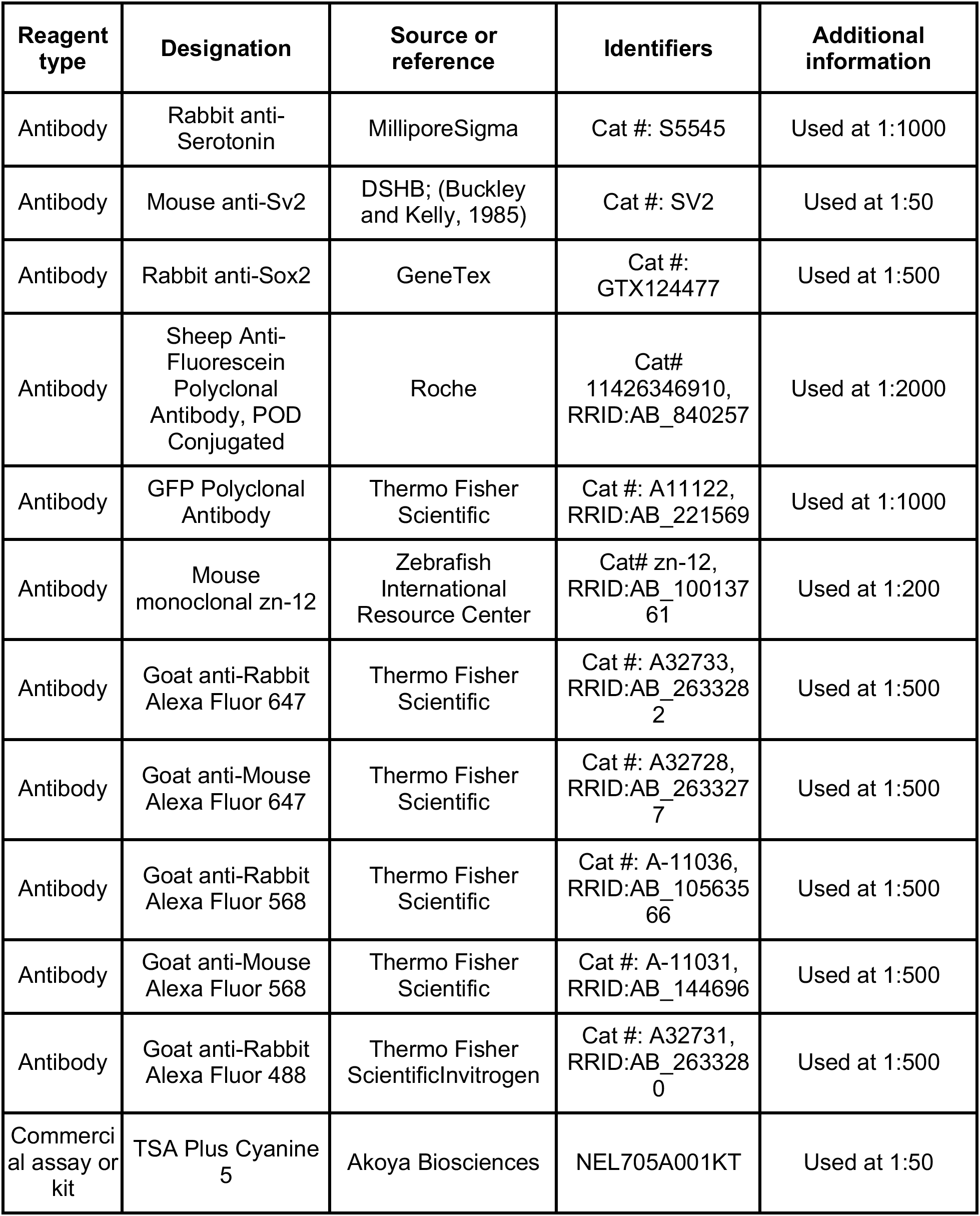
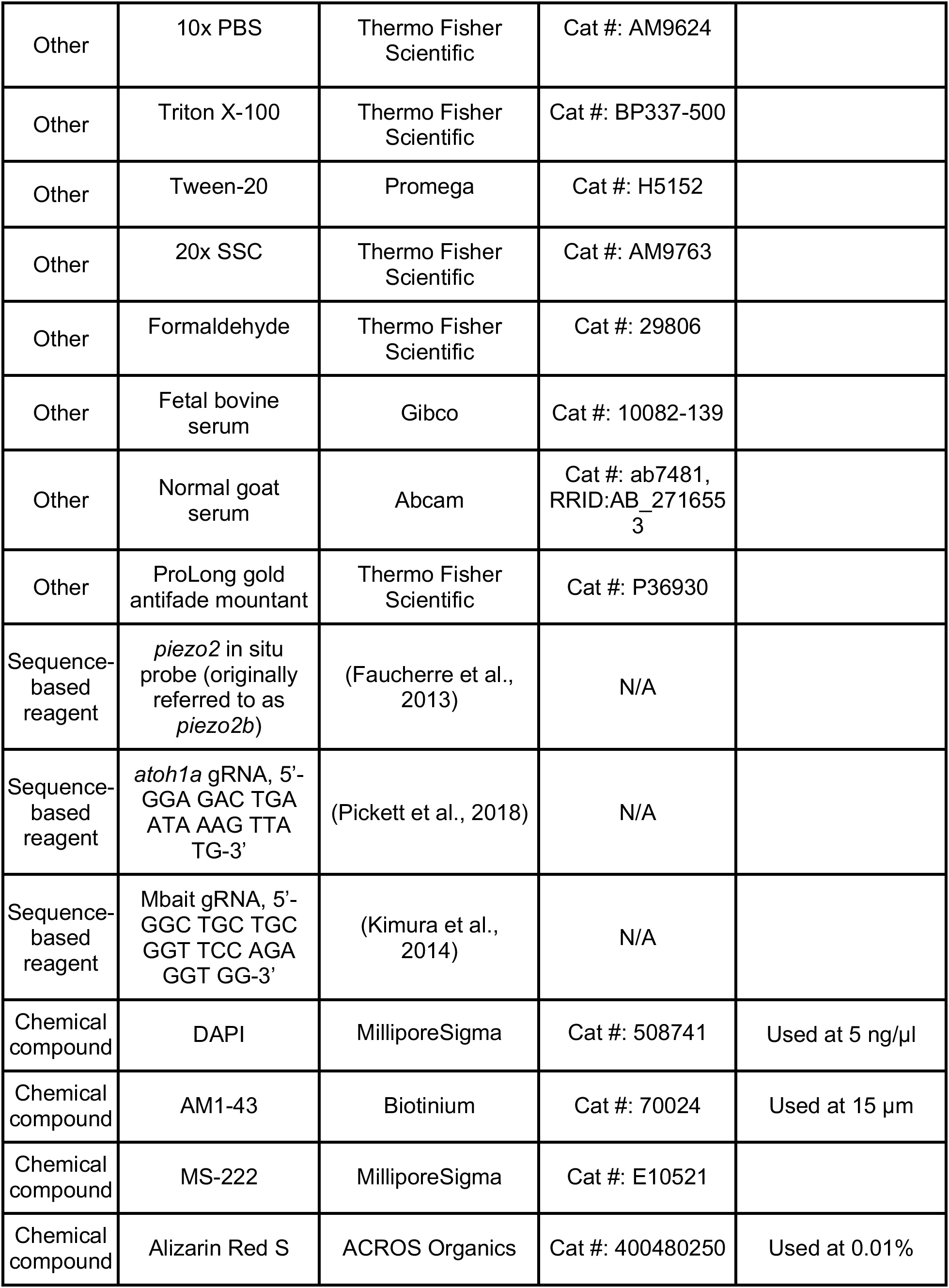
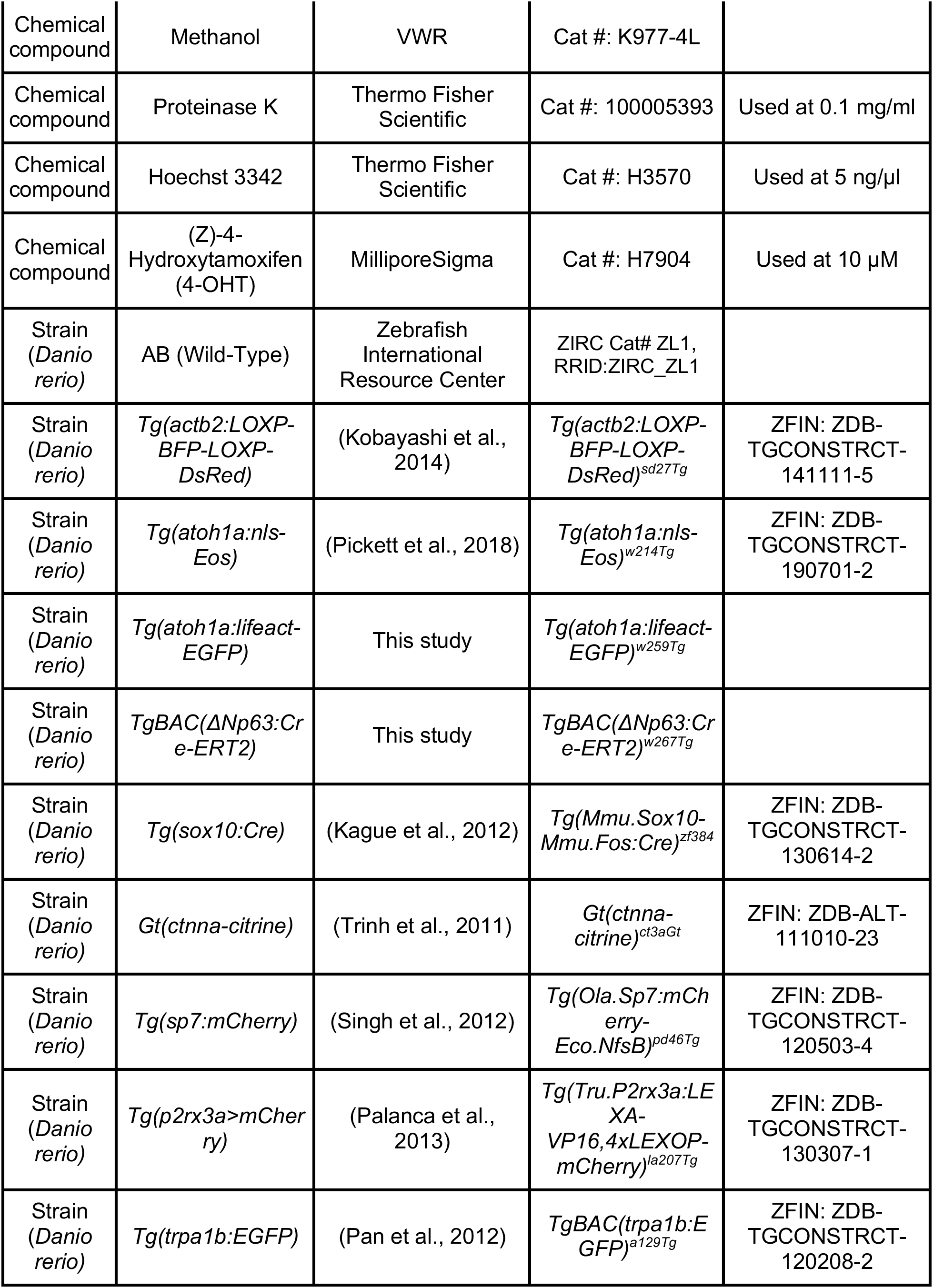
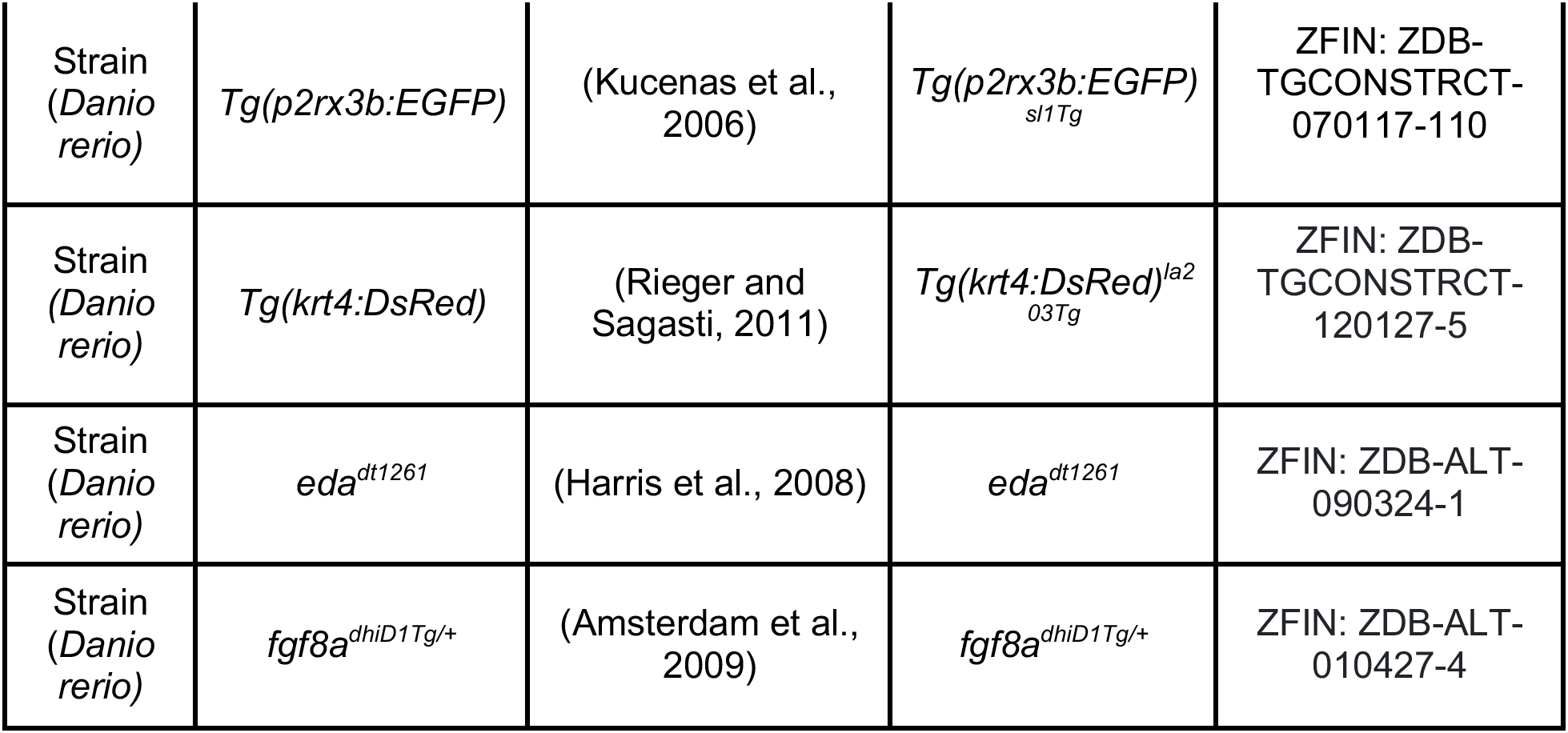

### Animals

#### Zebrafish

Zebrafish were housed at 26-27°C on a 14/10 h light cycle. See Key Resource Table for strains used in this study. Animals of either gender were used. Zebrafish were staged according to standard length (SL) (Parichy et al., 2009). SL of fish was measured using the IC Measure software (The Imaging Source) on images captured on a Stemi 508 stereoscope (Zeiss) equipped with a DFK 33UX264 camera (The Imaging Source). All zebrafish experiments were approved by the Institutional Animal Care and Use Committee at the University of Washington (Protocol: #4439-01).

#### Creation of Tg(atoh1a:Lifeact-EGFP)

*Tg(atoh1a:Lifeact-EGFP)^w259Tg^* was generated by CRISPR-mediated knock-in as previously described (Kimura et al., 2014). A donor plasmid containing the Mbait, minimal *hsp70l* promoter, *Lifeact-EGFP,* and *bgh poly(A)* sequences was created using Gibson assembly. The insertion was targeted 372 bp upstream of the endogenous *atoh1a* coding sequence using a previously published guide RNA (gRNA) (Pickett et al., 2018). The *Mbait-hsp70l-Lifeact-EGFP* plasmid, Mbait and *atoh1a* gRNAs, and Cas9 protein were prepared and injected into single cell embryos of the AB strain as previously described (Thomas and Raible, 2019). Larvae were screened for Lifeact-EGFP expression at 3 dpf and raised to adulthood. A founder adult was identified and outcrossed to generate a stable transgenic line.

#### Creation of TgBAC(ΔNp63:Cre-ERT2) and induction with 4-OHT

The *ΔNp63:EGFP-2xFYVE* bacterial artificial chromosome (BAC) was created by modifying the previously generated BAC DKEY-263P13-iTol2-amp (Rasmussen et al., 2015). The predicted *ΔNp63* start codon was replaced by a *Cre-ERT2-pA-KanR* cassette that contained a zebrafish codon-optimized *Cre-ERT2* (Kesavan et al., 2018) using a previously described protocol (Suster et al., 2011). *TgBAC(ΔNp63:Cre-ERT2)^w267Tg^* was created by injecting *tol2* mRNA, which was transcribed from pCS2-zT2TP (Suster et al., 2011), and BAC DNA into one-cell stage embryos and screening adults for germline transmission. To activate Cre-ERT2, 1 dpf embryos were treated with 10 μM 4-OHT for 24 h. 4-OHT was prepared as described (Felker et al., 2016).

#### Mutant identification and analysis

*eda* mutants and siblings were sorted by visible phenotype starting at 7 mm SL. Mutants were grown separately from siblings. *fgf8a^dhiD1Tg/+^* fish were identified based on altered scale patterning and/or pigmentation (Kawakami et al., 2000).

### Imaging and photoconversion

#### Electron microscopy

Isolated scales were prepared for TEM as described (Sire et al., 1997b), with the following modifications: after dehydration, scales were treated with propylene oxide (PO), infiltrated with PO:Eponate 12, and embedded in Eponate 12. Semithin sections (0.2 μm) stained with toluidine blue were used for orientation. Thin sections (50 nm) were placed on Formvar coated copper slot grids, stained with saturated uranyl acetate and Reynolds’ lead citrate, and examined on a JEOL 100CX at 60 kV or a Philips CM100 at 80 kV.

#### Confocal image acquisition

Confocal z-stacks were collected using a A1R MP+ confocal scanhead mounted on an Ni-E upright microscope (Nikon) using a 16× water dipping objective (NA 0.8) for live imaging or 40× oil immersion objective (NA 1.3) for fixed image acquisition. Images acquired in resonant scanning mode were post-processed using the denoise.ai function in NIS-Elements (Nikon). For live imaging, zebrafish were anesthetized in a solution of 0.006-0.012% MS-222 in system water for 5 min. Anesthetized fish were mounted in a custom imaging chamber, partially embedded in 1% agarose and covered with tricaine solution. For Supplemental Video 1, a FLUOVIEW FV3000 scanning confocal microscope (Olympus) equipped with a 100× objective (NA 1.49) was used to collect a z-stack and 3D rendered with Imaris (Bitplane).

#### Whole animal photoconversion

Prior to imaging, *Tg(atoh1a:nls-Eos)* zebrafish were exposed to light from a UV LED flashlight (McDoer) for 15 min in a reflective chamber constructed from a styrofoam box lined with aluminum foil. A similar lateral region of the trunk was imaged over subsequent days identified by approximate body position below the dorsal fin and relative to underlying pigment stripes.

#### Regional photoconversion

After anesthetization and mounting as described above, the *Tg(atoh1a:nls-Eos)* reporter was photoconverted using the stimulation program of NIS-Elements with the 405 nm laser at 14-18% power for 30-45 s within a 500X500 pixel ROI with an area of 67055 µm^2^. The same lateral region of the trunk was imaged over subsequent days identified by body position under the dorsal fin, position relative to underlying pigment stripes, and presence of photoconverted cells.

### Staining

#### Alizarin Red S Staining

To visualize mineralized bone, live animals were stained for 15 min in a solution of 0.01% (wt/vol) Alizarin Red S dissolved in system water, and subsequently rinsed 3×5 min in system water prior to imaging as described (Bensimon-Brito et al., 2016).

#### Antibody staining

Zebrafish were anesthetized in a solution of 0.012% MS-222 in system water for 5 min. Using metal forceps, up to 10 scales were removed from the lateral side of the trunk in the region below the dorsal fin. Scales were fixed in 4% PFA/PBS at 4°C overnight. Scales were washed 4×5 min in 1x PBS + 0.3% triton-X (PBST) at room temperature and then blocked for 1.5 h with PBST containing 5% normal goat serum. Incubation with primary antibodies occurred at 4°C overnight, followed by 4×15 min washes in PBST. Scales were incubated in appropriate secondary antibodies for 2 h at room temperature and washed 4×15 min in PBST. To label nuclei, scales were incubated with DAPI for 5 min at 4°C and washed in PBST 4×5 min at room temperature. Scales were mounted between a microscope slide and coverslip in Prolong gold. All steps were performed on a rotating platform.

#### AM1-43 staining

Scales were removed from adult *Tg(atoh1a:nls-Eos)* zebrafish as described above, and placed into the center of a petri dish. 1 mL of L-15 media was added to the dish containing newly plucked scales no longer than 2 min after the scales had been removed. 1.5 µl of 10 mM AM1-43 was added to the dish for a final concentration of 15 µM AM1-43. Scales were incubated for 5 min in this solution to allow for incorporation. Prior to confocal imaging, regional photoconversion of nls-Eos was carried out as described above.

#### Fluorescent in situ hybridization (FISH)

*piezo2 a*ntisense RNA was transcribed *in vitro* from a previously generated plasmid (Faucherre et al., 2013) using SP6 and fluorescein-dUTP. The FISH protocol for adult zebrafish scales was previously described (Lin et al., 2019). Briefly, scales from *Tg(atoh1a:Lifeact-EGFP)* adults were plucked and fixed in 4% PFA overnight at 4°C then washed three times with 1x PBS + 0.1% Tween-20 (PBSTw). Scales were dehydrated in sequential washes of 75% PBSTw:25% methanol (MeOH), 50% PBSTw:50% MeOH, 25%PBSTw:75%MeOH, then placed in 100% MeOH at −20°C overnight. Scales were rehydrated in sequential washes of 25% PBSTw:75% MeOH, 50% PBSTw:50% MeOH, 75%PBSTw:25% MeOH, then washed 3x in PBSTw. Scales were treated with 0.1mg/ml proteinase K for 5 min, then re-fixed in 4% PFA for 20 min. Scales were washed once in PBSTw, washed once in 50% PBSTw:50% hybridization buffer, then incubated in hybridization buffer for 2 h at 65°C. Scales were incubated in hybridization buffer with probe (∼1 ng/µl) overnight at 65°C. Scales were sequentially washed at 65°C in 75% hybridization buffer:25% 2xSSC + 0.1%Tween20 (SSCT), 50% hybridization buffer:50% 2x SSCT, 25% hybridization buffer:75% 2xSSCT, followed by 3 washes at room temperature in 2x SSCT, followed by 3 washes in 0.2x SSCT. Scales were then washed 3x in 1x PBS + 0.2% Triton X-100 (PBSTr), then blocked for 2 h in PBSTr + 5% FBS. Scales were incubated in blocking buffer with anti-fluorescein POD fragments (1:2000) overnight at 4°C. Scales were washed 6x in PBSTr, followed by staining with TSA Plus Cyanine 5 (1:50 dilution) for 10 min.

Following FISH, scales were incubated in PBSTr + 10%NGS for 2 h at room temperature. Scales were stained with an anti-GFP antibody (1:1000) in PBSTr + 10% NGS overnight at 4°C. Scales were washed in PBSTr, then incubated in secondary antibodies (1:200) for 2 h at room temperature. Scales were washed in PBSTr, stained with Hoechst (3.24 nM) for 10 min at room temperature, washed in PBSTr, mounted under coverslips in ProLong Gold, and imaged.

### Image analysis

#### Axon contact quantification

Innervation of *Tg(atoh1a:nls-Eos)-*expressing cells was scored using a custom ImageJ macro. A cell was scored as innervated if an axon passed within a sphere (created using the “3D project” function) centered around the nuclear center of mass that was 10% larger than the maximum nuclear diameter. In some cases, the *Tg(atoh1a:nls-Eos)* reporter was photoconverted prior to image acquisition as described above.

#### Cell density analysis

Maximum intensity projections of confocal z-stacks were converted to 8-bit images and thresholded in ImageJ. Cell density was quantified using the “Analyze particles” function of ImageJ. For low magnification quantification of MC cell density across the trunk of *fgf8a^dhiD1Tg/+^*and siblings, tiled images were collected that included multiple scales per region. Cell density was quantified as described above using ImageJ. For high magnification cell density quantification in the epidermis directly above scales, a small region centered in the epidermis of each full scale in view and positioned based on scale lobe was quantified. For scales with multiple lobes, a density measurement was collected from the center of each lobe and averaged.

#### Statistical analysis

Statistical tests used are listed in individual figure legends. Plots were created using R or Python.

## ACKNOWLEDGEMENTS

We thank the LSB Aquatics staff for animal care; Wai Pang Chan and Marianne Cilluffo for TEM support; the labs of Ajay Dhaka, Jacqueline Lees, and Alvaro Sagasti for sharing zebrafish stocks; the lab of Chris Joplin for sharing the *piezo2* plasmid. The authors are grateful to all members of the Rasmussen lab for discussion, technical assistance, and support.

## CONFLICT OF INTERESTS

The authors declare that they have no conflict of interest.

## FUNDING

This work was funded in part by a Postdoctoral Fellowship (#2011008) from the National Science Foundation to TLB, a Graduate Research Fellowship (DGE-2140004) from the National Science Foundation to EWC, R01HD107108 from the Eunice Kennedy Shriver National Institute of Child Health and Human Development to JPR, A153025 from the University of Washington Research Royalty Fund to JPR, and a New Investigator Award from the University of Washington/Fred Hutchinson Cancer Research Center Cancer Consortium, which is supported by the NIH/NCI Cancer Center Support Grant P30 CA015704, to JPR. JPR is a Washington Research Foundation Distinguished Investigator.

---

**Figure 1—figure supplement 1.**
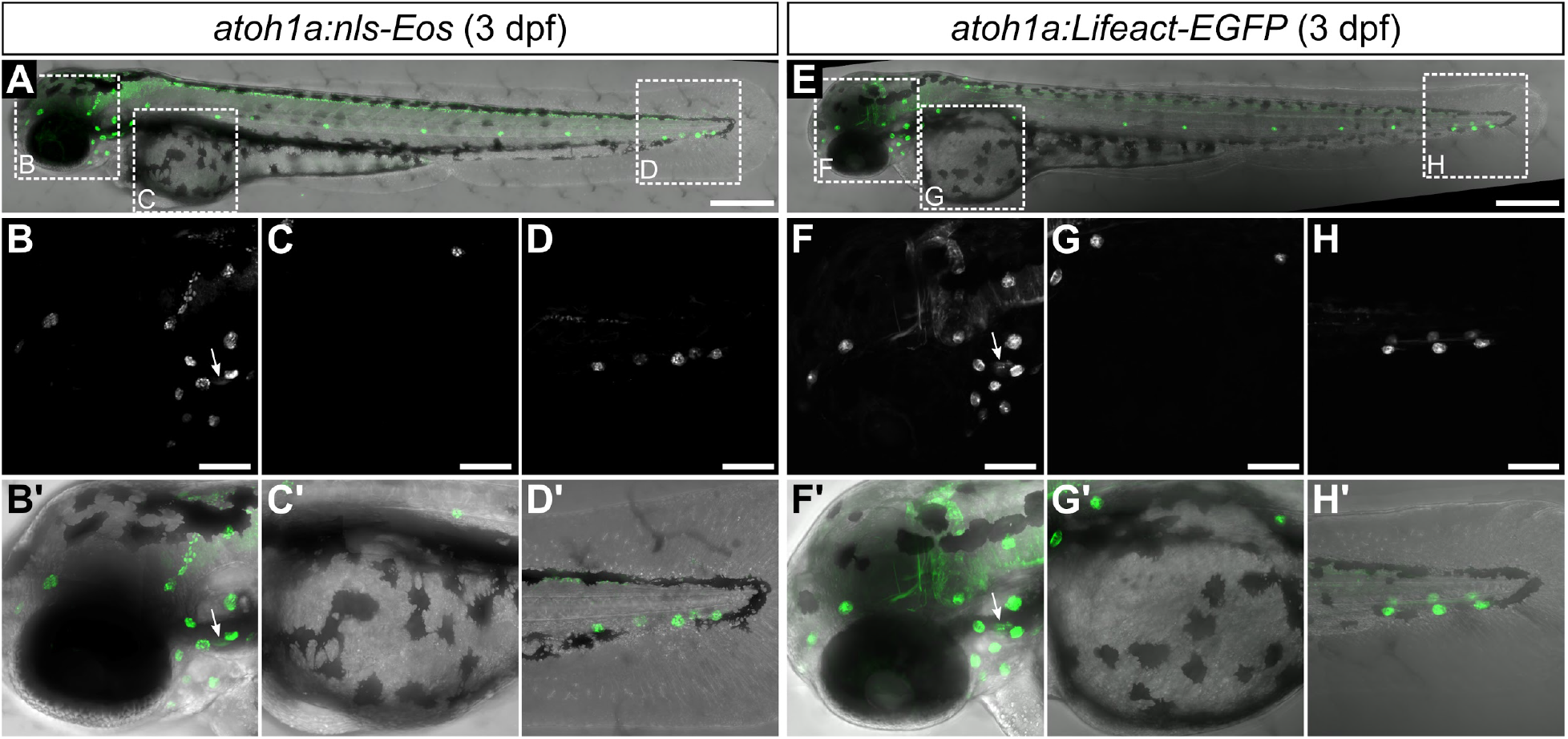
Characterization of *atoh1a* reporter transgenes in larvae. Confocal micrographs of 3 dpf larvae expressing the indicated transgenes. Dotted boxes in A and E indicate areas of magnification for panels below. Arrows indicate expression by hair cells of the inner ear. Scale bars: 300 μm (A,E) and 100 μm (B-D,F-H).

**Figure 1—figure supplement 2.**
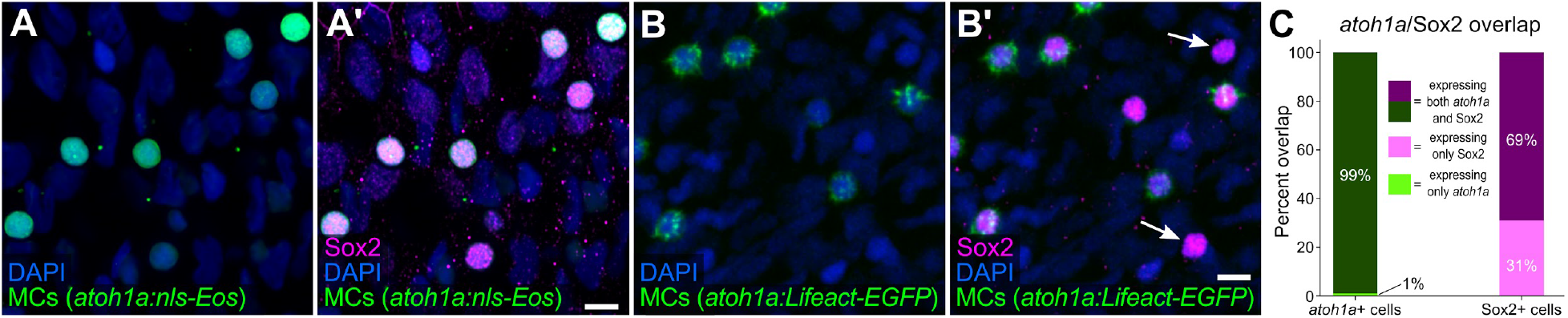
MCs in the adult epidermis express Sox2. **(A-B’)** Lateral confocal micrographs of the scale epidermis showing anti-Sox2 immunostaining of MCs labeled by either *Tg(atoh1a:nls-Eos)* (A,A’) or *Tg(atoh1a:Lifeact-EGFP)* (B,B’). Arrows indicate examples of Sox2+/*atoh1a-* cells. DAPI labels epidermal nuclei. **(C)** Quantification of the overlap between *atoh1a*+ MCs and Sox2 immunostaining. 99% of *atoh1a:nls-Eos+* MCs expressed Sox2 (769/774 cells from *N*=5 fish). Scale bars: 5 μm.

**Figure 4—figure supplement 1.**
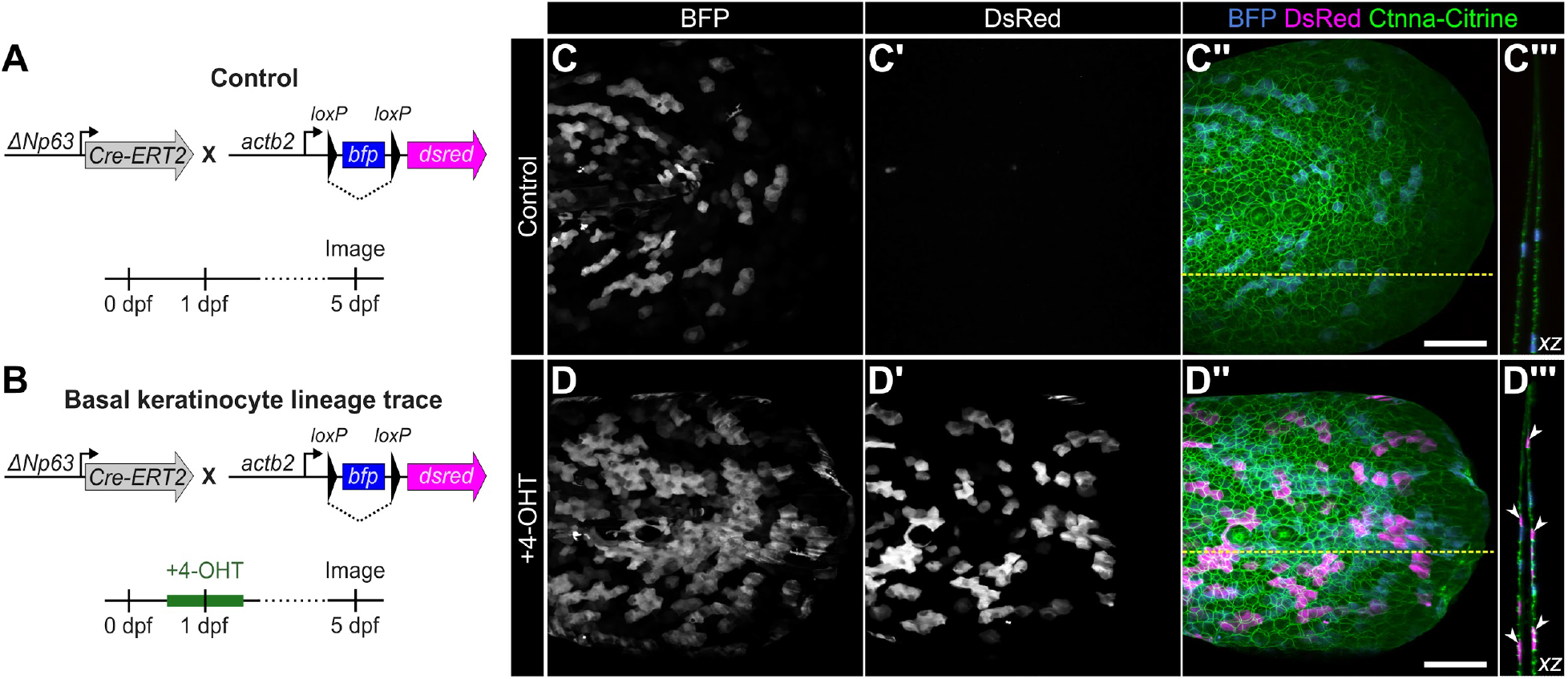
Validation of basal keratinocyte lineage tracing strategy. **(A,B)** Schematics of the experimental design. **(C-C’’,D-D’’)** Lateral confocal micrographs of the caudal fin of *TgBAC(ΔNp63:Cre-ERT2); Tg(actb2:LOXP-BFP-LOXP-DsRed)* larvae treated as indicated. *Gt(Ctnna-Citrine)* labels keratinocyte membranes. **(C’’’,D’’’)** Reconstructed cross sections along the dashed yellow line in C’’ or D’’. Arrowheads indicate examples of basal keratinocytes that have undergone Cre recombination as evidenced by DsRed expression. Scale bar: 100 μm.

**Figure 8—figure supplement 1.**
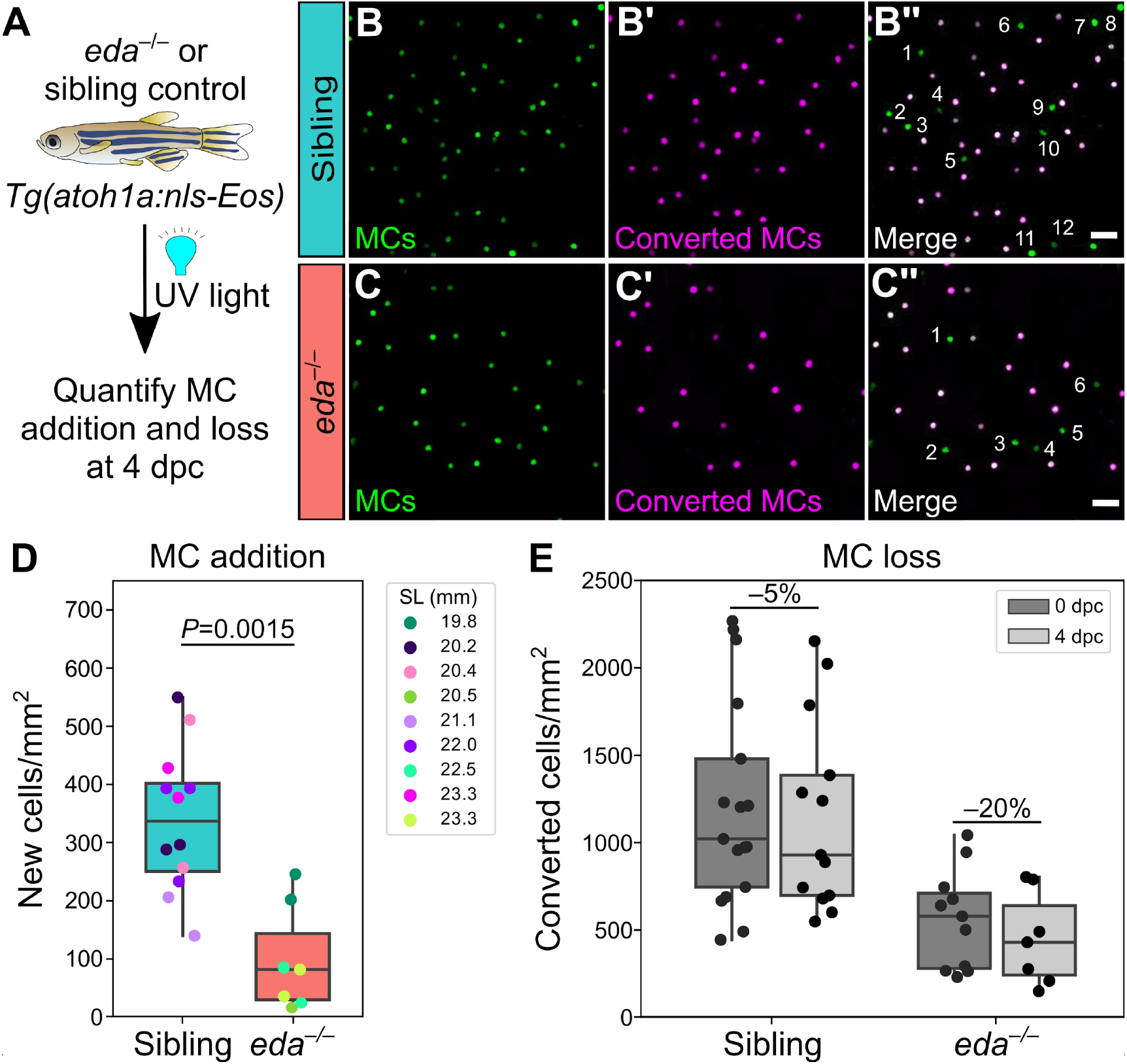
*eda* mutants exhibit decreased MC addition and increased MC loss. **(A)** Schematic of experimental approach. Following whole animal photoconversion, densities of converted and non-converted MCs were quantified at 4 dpc. **(B-C’’)** Representative lateral confocal micrographs of MCs at 4 dpc in adults of the indicated genotypes. Numbers label newly added cells, distinguishable by the absence of photoconverted nls-Eos (magenta). **(D)** Boxplots of MC addition at 4 dpc in the indicated genotypes. 1-3 independent regions were analyzed per animal. *eda* mutants show a significantly lower rate of MC cell addition (Mann-Whitney test). **(E)** Boxplots of photoconverted MC density in animals of the indicated genotypes. Average percentage cell density loss between 0 and 4 dpc is listed above the boxplots for each genotype. Scale bars: 20 μm (B,C).

**Figure 9—figure supplement 1.**
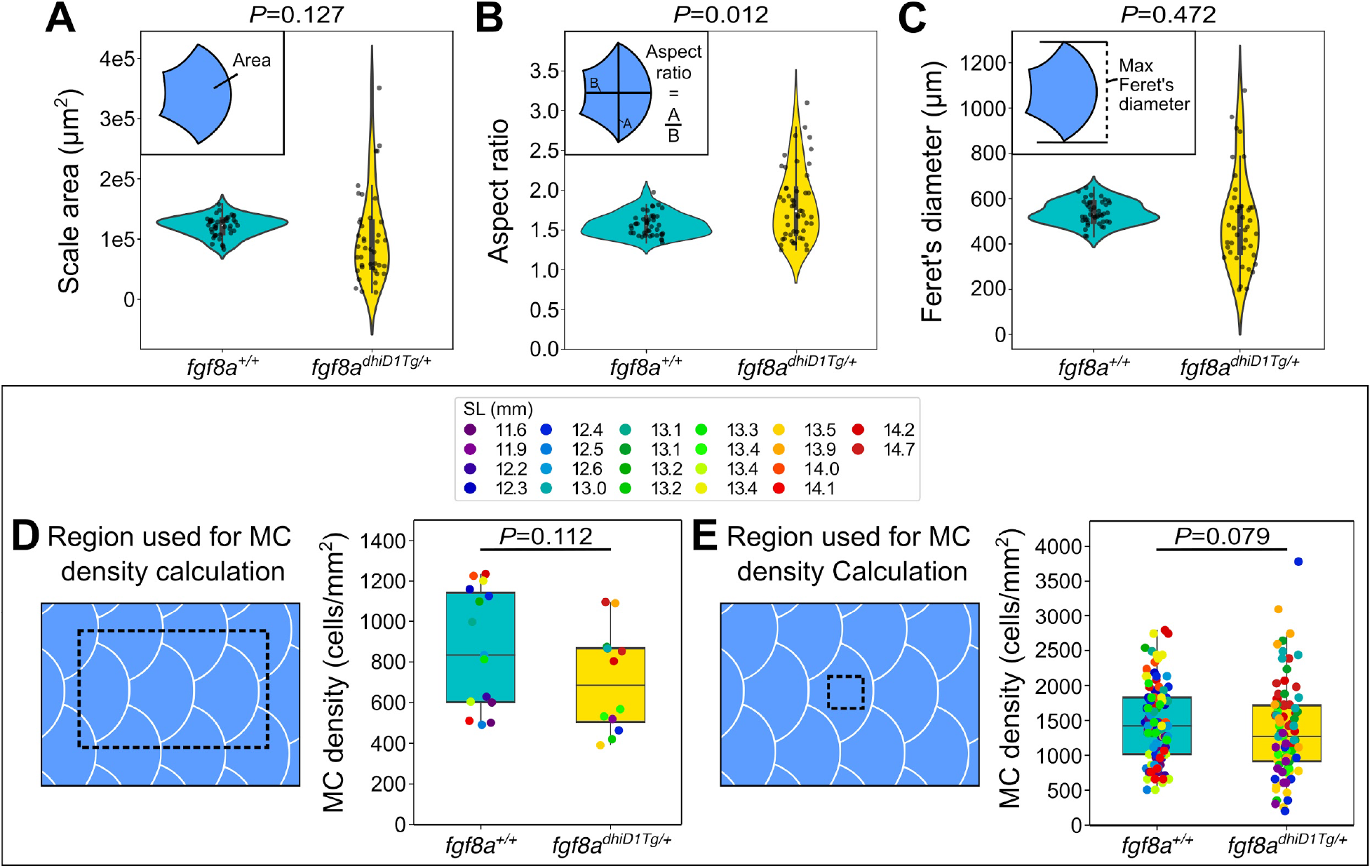
*fgf8a^dhiD1Tg/+^* juveniles show altered dermal appendage size and shape, but not MC density. **(A-C)** Violin plots of scale area, aspect ratio, and Feret’s diameter from juveniles (11.6-14.7 mm SL) of the indicated genotypes. *P*-values (Mann-Whitney test), listed above each plot, indicate a significant difference between the genotypes for the scale aspect ratio, but not scale area or Feret’s diameter. Data represent *n*=42 scales from *N*=13 fish (*fgf8a^+/+^*) and *n*=32 scales from *N*=9 fish (*fgf8a^dhiD1Tg/+^*). Insets illustrate the various measurements. **(D,E)** Boxplots of MC density across the trunk epidermis (D) or the epidermis directly above individual scales (E) as indicated by the dotted boxes in juveniles expressing a MC reporter (*Tg(atoh1a:nls-Eos)*). Dot colors represent animal SL as indicated in the legend. Total fish analyzed: *fgf8a^+/+^* (*N*=13); *fgf8a^dhiD1Tg/+^*(*N*=9). *P*-values (Mann-Whitney test) are listed above each plot.

## SUPPLEMENTAL VIDEO LEGENDS

### Supplemental Video 1

3D rotation of somatosensory axons (green) and photoconverted MCs (green and magenta) in the adult scale epidermis. Arrows indicate axonal varicosities in close proximity to MCs.

## Notes

### Competing Interest Statement

The authors have declared no competing interest.

### Summary of Updates

Supplemental files updated.

